# Context-specific genetic interaction mapping reveals combinatorial KRAS/FGFR dependence in pancreatic cancer

**DOI:** 10.64898/2026.09.24.754062

**Authors:** Richard Y. Ebright, Yao He, Alan G. Zhang, Ian C. McCabe, Chinmaya U. Joisa, Tanay Thakar, Jayu Jen, Dennie T. Frederick, Arianny Acosta, Guangyan Li, Elena Kuehner, Vivian Yang, Katherine E. Doherty, Natnael Yaregal, Ngan N. K. Van, Sandra A. Zarmer, Changfei Luan, Aurelia Dingley, Eiman Elwakeel, Yuen-Yi Tseng, Andrew J. Aguirre, James M. Cleary, Jen Jen Yeh, William R. Sellers

## Abstract

Oncogenic KRAS drives >90% of pancreatic ductal adenocarcinoma (PDAC), and the pan-RAS inhibitor daraxonrasib doubles overall survival, but adaptive and acquired resistance limit the depth and duration of responses. To systematically map combination vulnerabilities, we developed RAS+, a digenic CRISPR knockout library testing 10,296 pairwise perturbations across oncogenic signaling pathways, and screened 12 cell lines, revealing genotype- and lineage-specific interactions. In KRAS^mut^ PDAC, the most differentially effective gene pair was KRAS and FRS2, an adapter coupling FGFR signaling to RAS. Combined KRAS/pan-FGFR inhibition was synergistic and cytotoxic, eradicating tumor cells and overcoming resistance in vitro and deepening and prolonging regressions in vivo. Single-cell analysis identified cancer-associated fibroblasts (CAFs) as a primary source of FGF ligands in PDAC, and CAF conditioned media or individual FGFs promoted resistance. These findings define a stroma-to-tumor adaptive FGFR circuit limiting the effects of KRAS inhibition. Thus, combined KRAS/FGFR blockade may be a strategy to improve responses.

**STATEMENT OF SIGNIFICANCE:** The recent approval of daraxonrasib makes RAS inhibition standard-of-care in pancreatic cancer, but resistance inevitably develops. Unbiased digenic CRISPR screening identified FGFR signaling as a prominent resistance axis potently activated by stromal FGF ligands. The addition of an approved FGFR inhibitor to RAS inhibition eradicates tumor cells and prolongs response, offering an immediately actionable combination.

## INTRODUCTION

The RAS/MAPK and PI3K signaling pathways are commonly dysregulated in human cancer, with oncogenic mutations in either pathway or upstream receptor tyrosine kinases driving tumor cell proliferation and survival (1). Direct inhibition of pathway components has provided substantial clinical benefit to patients. Unfortunately, single-agent efficacy is often limited, with many inhibitors yielding modest response rates of relatively short duration due to the development of resistance. Preclinical and clinical studies have identified reactivation of MAPK and/or PI3K signaling and activation of alternative pathways such as YAP/TAZ/TEAD as causes of resistance. Thus, these pathways interact in clinically-impactful ways (2–4).

Such pathway interactions can be targeted to overcome resistance. For instance, in BRAF^V600E^ and KRAS^G12C^ colorectal cancer (CRC), BRAF- and KRAS-targeted therapies demonstrate modest monotherapy clinical efficacy due to adaptive EGFR signaling reactivating RAS/MAPK signaling (5–7). Concomitant EGFR inhibition blocks this adaptive resistance, leading to greater depth, duration, and frequency of clinical responses (8–11). Similarly, in EGFR^mut^ non-small cell lung cancer (NSCLC), MET activation leads to EGFR inhibitor resistance (12), which can be overcome by MET inhibition (13). Unfortunately, the broader development of efficacious targeted therapy combinations has been limited by the difficulty of identifying actionable genetic interactions and by the exacerbation of clinical toxicities observed when combining agents (14).

To address the first of these factors, we turned to combinatorial digenic CRISPR knockout screening. We previously developed digenic libraries using a dual-Cas9 system to evaluate genetic interactions between paralogous gene pairs, identifying DUSP4/DUSP6 and 14-3-3ε/ζ paralog dependence in NRAS^mut^ melanoma (15). Subsequently, a dual-tracer system was used to develop a larger paralog targeting library (PARADIGM) which has been screened across 283 cancer cell lines spanning 19 lineages (16). However, data from the PARADIGM screens demonstrated that dual loss of paralogs (e.g., CDK4/CDK6) was often broadly pan-lethal and *less* tumor-selective than individual paralog loss; thus, refining this approach to discover non-paralogous digenic dependencies is of high interest.

Here, we developed a new digenic CRISPR knockout library. The **“RAS+”** library assesses all pairwise knockouts of 144 genes in the RAS/MAPK, PI3K, and YAP/TAZ pathways, as well as upstream RTKs, RTK adapters, and downstream cell cycle mediators. Each of these pathways is known to contribute to KRAS inhibitor resistance (17,18). In total, 10,296 combination knockouts are assessed simultaneously, along with the corresponding single gene knockouts.

The RAS+ library was screened across 12 cancer cell lines. Gene pair knockouts exhibiting synergistic cell growth inhibition were identified in each cell line, as well as in genomically-defined subsets of the screened lines. Notably, the genetic interaction maps of the subsets were strikingly different, with digenic dependencies in KRAS^mut^ lines centered on pairs containing KRAS, while digenic dependencies in BRAF^mut^ or PIK3CA^mut^ lines demonstrated much greater variation.

Interestingly, the RAS+ screens revealed a specific combinatorial knockout effect of KRAS and FGFR pathway members in KRAS^mut^ pancreatic cancer (PDAC). KRAS mutations are the oncogenic drivers in >90% of PDAC (19). However, the role of FGFR signaling in PDAC is much less clearly defined, though FGFR copy number gains or fusions are seen in KRAS^WT^ PDAC (20–22). We validated the genetic interaction between KRAS and FGFR signaling *in vitro* and *in vivo* using both genetic and pharmacologic approaches. Mechanistically, we found that prolonged KRAS inhibition leads to adaptive FGFR pathway activation, and we identified cancer-associated fibroblasts (CAFs) as a key source of FGF ligands in the pancreatic tumor microenvironment (TME). These findings suggest that combined KRAS/FGFR inhibition may be effective in PDAC patients.

## RESULTS

### Generation of a novel digenic CRISPR knockout library for genetic interaction mapping

To map genetic interactions within and among oncogenic pathways, we developed a digenic CRISPR library targeting the RAS/MAPK, PI3K, and YAP/TAZ signaling pathways, including upstream RTKs and downstream cell cycle mediators. We selected 144 genes covering key pathway components. These genes include oncogenes (e.g., KRAS, BRAF, PIK3CA, EGFR, YAP1), adapter proteins (e.g., GRB2, PTPN11, FRS2), regulators of signaling (e.g., DUSP4, PEA15), tumor suppressors (e.g., PTEN, NF1), scaffolding proteins (e.g., KSR1, KSR2), and core cell cycle regulators (e.g., CDK2/4/6, CCND1-3, CCNE1-2) (**Fig. 1A, Table S1**). Pan-essential genes, defined as genes whose loss broadly impairs cellular fitness across diverse cell lineages, were excluded, as the strong lethality of individual knockouts would limit the ability to observe dual knockout synergy (23). The RAS+ library assesses all possible pairwise combinations of the 144 genes, yielding 10,296 gene pairs, as well as 145 pan-essential and 143 non-essential positive and negative control genes. We also paired guides for each individual gene with a control guide targeting the safe-harbor AAVS1 locus, providing internal controls for the effects of CRISPR cutting toxicity. The non-essential gene_AAVS1 guide pairs provided an empirical background distribution of fitness effects for screen normalization and quality assessment.

**Figure 1.**
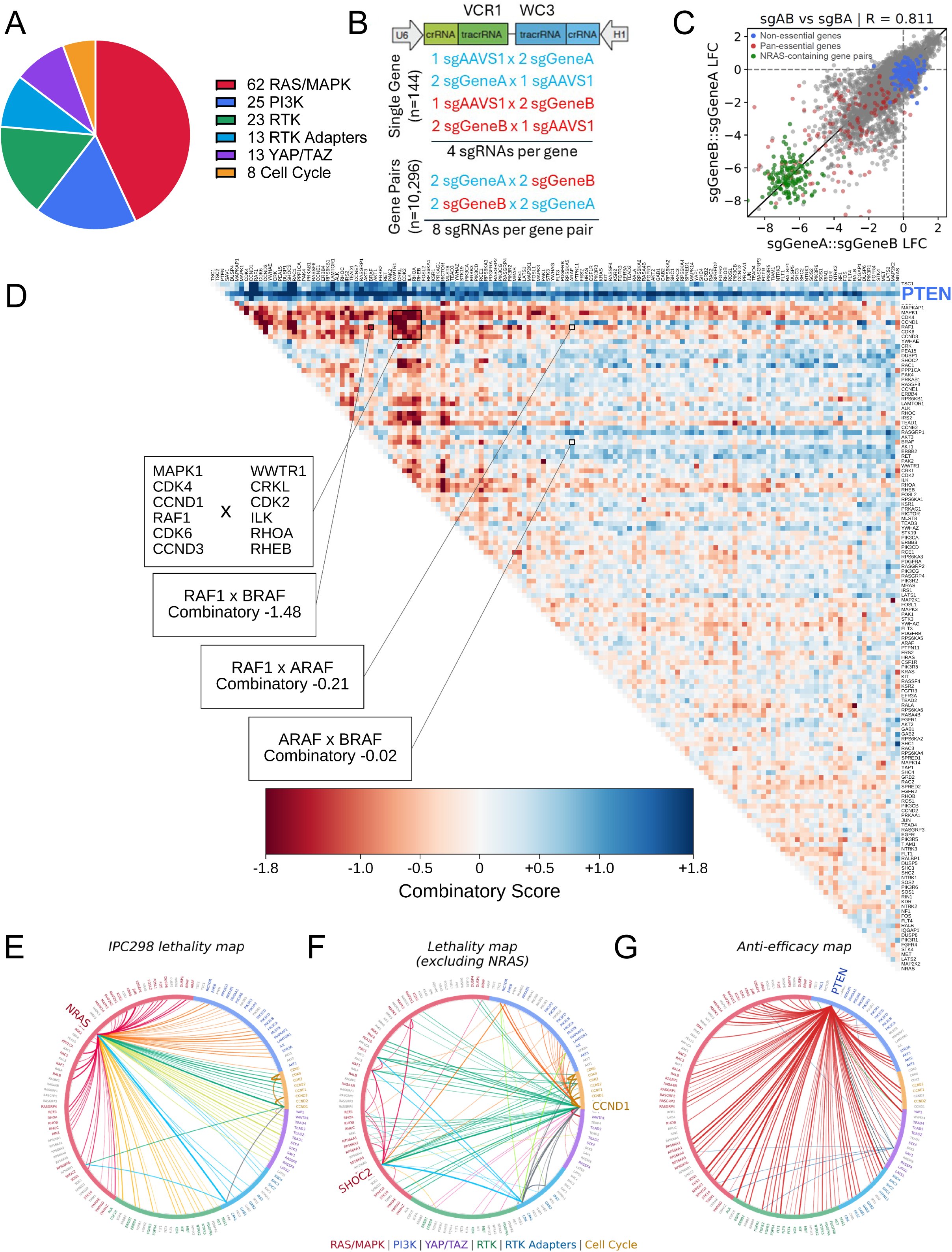
Development of the digenic RAS+ library. (A) Breakdown of 144 genes in the RAS/MAPK, PI3K, and YAP/TAZ signaling pathways, as well as upstream RTKs and adapters and downstream cell cycle genes. (B) Design of the sgRNA library for single genes and gene pairs. Single genes are targeted by 4 different sgRNAs, 2 driven by the U6 promoter and 2 driven by the H1 promoter. Gene pairs are targeted by 8 different sgRNA pairs. In total, 10,296 distinct gene pairs are assessed. (C) Correlation between average log fold change (LFC) of dual knockouts in the NRAS^mut^ melanoma cell line IPC298 when sgRNAs are swapped between U6 and H1 promoter. (D) Combinatory score genetic interaction map comprising all 144 genes in the RAS+ library, with genes ordered by hierarchical clustering of Combinatory score profiles. Red indicates synergistic interactions (negative Combinatory scores), and blue indicates antisynergistic interactions (positive Combinatory scores). Callouts as discussed in the text. (E-F) Lethality maps showing the 100 most lethal gene pairs in IPC298 (E) and the 100 most lethal gene pairs in IPC298 which do not include NRAS (F). Chords are colored by which gene categories are connected by each chord, and greater width indicates greater lethality. (G) Growth advantage map showing the 100 least lethal gene pairs in IPC298. Chords are colored by which tumor suppressor(s) are connected by each chord (PTEN: red; TSC1: green; SAV1: blue; multiple: gray), and greater width indicates greater growth advantage.

Building on our prior work (15,16), the RAS+ library uses a dual-promoter system (U6, H1) expressing two sgRNAs with alternative *S. pyogenes* Cas9 tracrRNAs that yield efficient and balanced dual knockouts. To reduce knockout variation, each gene is targeted with four sgRNAs combined with an AAVS1-targeting guide, and each gene pair is targeted with eight sgRNA pairs (**Fig. 1B**). To normalize differential effects between promoters, two guides are expressed from each promoter. To maximize the probability of selecting high performing guides, we leveraged our prior dual-guide screens (15,16) and the DepMap genome-wide Avana screens to find effective guides (24). Random forest models were used to assign agreement scores to guides based on consistent behavior across cell lines, allowing us to prioritize guides most likely to effectively knockout the gene of interest (**Fig. S1A; Methods**). Based on this modeling, we selected guides directly from the PARADIGM or Avana libraries predicted to be efficacious, covering 94% of guides in the library. The remaining 6% of guides were designed *de novo* via crisprDesign (25). In total, the RAS+ library comprises 84,200 constructs targeting 10,296 gene pairs (**Table S2**).

Compared to our prior digenic libraries and others, the RAS+ library has a more uniform plasmid DNA (pDNA) sgRNA representation, with 90% of pDNA reads contributed by 83% of sgRNAs (**Fig. S1B**) (15,26–29). We first screened the RAS+ library in IPC298, an NRAS^mut^ melanoma line which we previously screened with two digenic CRISPR libraries (**Methods**) (15,16). The RAS+ library demonstrated good screening quality with the receiver operating characteristic (ROC)-area under the curve (AUC) and null-normalized mean difference (NNMD) score (-10.2) demonstrating robust performance of positive and negative controls comparable to our prior digenic libraries (**Fig. S1C**). Promoter effects between the U6 and H1 promoters were minimal (**Fig. 1C**). We calculated naïve log2-fold change (LFC) values from the guide pair abundance in screen output compared to the pDNA, and guide pair LFCs for each gene pair were aggregated into gene pair LFC scores (**Methods**). LFC scores for gene pairs shared between the RAS+ library and the Avana or PARADIGM library were comparable (**Fig. S1D-E**). Finally, RAS+ screening at 500x or 250x coverage yielded comparable results, highlighting the uniformity of guide distribution in the pDNA enabling equivalent results at lower cell coverage (**Fig. S1F**).

### The RAS+ library identifies highly lethal and synergistic gene pairs in IPC2G8

IPC298 screening data were analyzed to identify genetic interactions between gene pairs. We calculated a Combinatory score, measuring the difference between the observed double knockout effect and the more growth inhibitory of either single gene knockout. Combinatory scores <-0.5 imply a negative/synergistic genetic interaction and >0.5 imply a positive/antagonistic genetic interaction (**Methods**). Unsupervised hierarchical clustering of Combinatory scores across the library yielded a genetic interaction map of RAS+ genes (**Fig. 1D**). Most gene pairs had Combinatory scores <|0.5|, indicating little to no genetic interaction. However, specific gene pair clusters exhibited strong, negative genetic interactions. One such cluster consisted of key MAPK/Cell Cycle genes (RAF1, MAPK1, CCND1, CCND3, CDK4, CDK6) paired with canonical nodes outside of the RAS/MAPK pathway (WWTR1, CRKL, ILK, RHOA, RHEB). IPC298 is an NRAS^mut^ line with strong dependency on MAPK signaling for proliferation; thus, some of the strongest genetic interactions are derived from MAPK pathway inhibition in combination with loss-of-function perturbations in other pathways. We also identified differences in paralogous gene pair behavior, such as the strong genetic interaction of BRAF_RAF1 (Combinatory: -1.48) but a lack of genetic interaction for ARAF_BRAF (Combinatory: -0.02) or ARAF_RAF1 (Combinatory: -0.21), despite mRNA expression of all three paralogs. Conversely, knockout of the tumor suppressor genes PTEN, SAV1, TSC1, and TSC2 all demonstrated antagonistic effects when combined with many other genes in the library. Interestingly, PTEN disruption had the strongest antagonistic effects, with Combinatory scores >0.5 when paired with inactivation of 99% (142/143) of other genes.

We next generated overview maps assessing and ranking growth inhibition by gene pairs. In IPC298, gene pairs including NRAS dominated this landscape, comprising 93 of the 100 most effective pairs, suggesting that few gene pairs are as lethal as gene pairs that include NRAS (**Fig. 1E**). Interestingly, SHOC2 was one of the few nodes distinct from NRAS, consistent with the recent observation that NRAS^mut^ cancers are SHOC2-dependent (30,31). Assessing the 100 most effective NRAS-independent pairs further highlights SHOC2 as a dependency node, as well as downstream cell cycle genes, particularly CCND1 (**Fig. 1F**). In contrast, the 100 most growth-promoting gene pairs were dominated by tumor suppressors, with PTEN pairs consistently demonstrating the greatest growth promotion (**Fig. 1G**).

### The RAS+ library identifies broadly recurrent genetic interactions

We screened the RAS+ library in 11 additional cancer lines of gastrointestinal origin (**Fig. S2A**). All screens demonstrated robust separation of control guides targeting pan- and non-essential genes, reflected in NNMD scores <-2.0 (**Fig. S2B**). LFC and Combinatory scores for each screen revealed distinct patterns of lethality and synergy across the lines, including among lines with overlapping genetic or lineage backgrounds (**Figs. S3-4; Tables S3-4**). As previously reported, paralog pairs were among the most significant genetic interactions (15,32); however, we also observed highly significant, non-paralogous genetic interactions present in individual lines (**Fig. 2A**). For instance, HCT15 is a KRAS^mut^/PIK3CA^mut^ CRC line, and KRAS_PIK3CA was the second-most synergistic gene pair (p = 1.17*10^-8^). Data from the 12 screens are available via the DepMap Portal (www.depmap.org).

**Figure 2.**
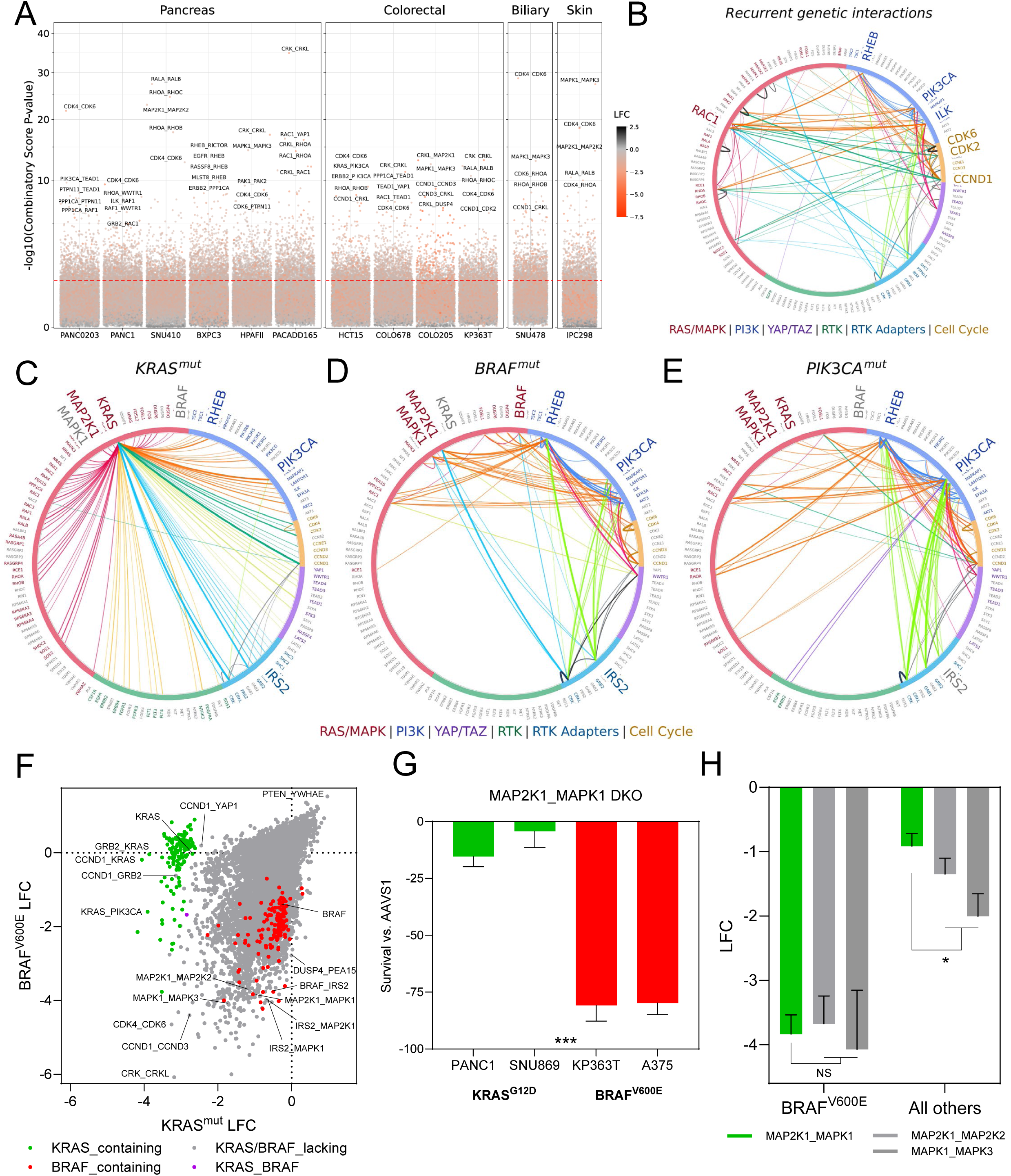
RAS+ library screening reveals lineage and genotype specific genetic interactions. (A) Manhattan plot of p values corresponding to Combinatory scores and color-coded by LFC. The top five most significant gene pairs are labeled for each cell line. (B) Chord diagram showing all gene pairs exhibiting genetic interactions (Combinatory score ≤ -0.5) in the majority of screened cell lines. Chords are colored by which gene categories are connected by each chord, except for paralog gene pairs, which are gray. Greater width indicates greater number of cell lines exhibiting genetic interactions. (C-E) Lethality maps showing the 100 most lethal gene pairs in KRAS^mut^ (C), BRAF^mut^ (D), or PIK3CA^mut^ (E) lines. Chords are colored by which gene categories are connected by each chord, and greater width indicates greater average lethality. (F) LFC plot in KRAS^mut^ versus BRAF^V600E^ lines. (G) Dual MAP2K1_MAPK1 knockout growth effects in BRAF^V600E^ versus KRAS^G12D^ lines. AAVS1 serves as a control for normalization, and viability measurements were made with CellTiter-Glo (CTG) readouts. (H) Dual MAP2K1_MAPK1, MAP2K1_MAP2K2, and MAPK1_MAPK3 data from the 12 RAS+ screens in BRAF^V600E^ versus all other screens. Data represent mean ± SD. *P* value calculated by two-tailed unpaired Student’s *t* test. ***: *P* < 0.001, *: *P* < 0.05, NS: *P* > 0.05.

To identify genetic interactions common across cancers, we assessed gene pairs with Combinatory score ≤-0.5 in more than half of our screens. This analysis identified 123 gene pairs, representing 1.2% of gene pairs assessed by the library (**Fig. 2B**). Given the limited diversity of lineages in these screens, we refer to these as “broadly recurrent” genetic interactions rather than pan-lethal. These broadly recurrent genetic interactions spanned a variety of functions, and most (70%; 86/123) were between genes in different pathways. Notably, there were no broadly recurrent genetic interactions that included RTKs, suggesting that genetic interactions including RTKs are restricted by genetic and/or lineage context. Of the remaining within-pathway genetic interactions, 32% (12/37) were between paralogs, representing some of the most recurrent genetic interactions across lines, including CDK4_CDK6 (12/12 lines), PAK1_PAK2 (11/12 lines), and RALA_RALB (11/12 lines). Other genes appeared to serve as genetic interaction hubs, forming interactions with many partners: genes involved in ≥10 recurrent genetic interactions included those involved with cell cycle progression (CCND1, CDK6, CDK2), integrins/motility (ILK, RAC1), and PI3K/MTOR signaling (PIK3CA, RHEB).

### The RAS+ library identifies lethal combinations across different oncogenic genetic backgrounds

We next assessed lethality across lines with shared genetic background, creating lethality maps for KRAS^mut^, BRAF^mut^, or PIK3CA^mut^ contexts. Like the NRAS^mut^ map (**Fig. 1E**), the KRAS^mut^ map converges on KRAS, with KRAS-containing gene pairs comprising 92 of the 100 most effective pairs (**Fig. 2C**). In contrast, BRAF^mut^ and PIK3CA^mut^ lethality maps were less BRAF- and PIK3CA-centric. In BRAF^mut^ lines, interaction nodes in addition to BRAF included MAP2K1 (MEK1), MAPK1 (ERK2), and IRS2 (**Fig. 2D**). Notably, IRS2, but not its paralog IRS1, was a selective combination partner with multiple MAPK pathway members: BRAF_IRS2, MAP2K1_IRS2, and MAPK1_IRS2 were all in the 100 most lethal pairs. IRS2 is a signaling adapter for insulin/IGF receptors and other RTKs, and ERK-dependent phosphorylation of IRS2 inhibits signaling (33,34). These findings suggest that MAPK suppression in BRAF^mut^ lines exposes an IRS2-specific adapter dependency, consistent with loss of ERK-dependent feedback inhibition (35). In PIK3CA^mut^ lines, PIK3CA and RHEB were the most common interaction nodes; both demonstrated many interactions with RTK adapters, suggesting a convergence of signaling from multiple upstream RTKs contributing to growth in the setting of pathway suppression (**Fig. 2E**). Intriguingly, RHEB is an activator of the mTORC1 but not mTORC2 complex (36), and mTORC2-specific complex members such as RICTOR and MAPKAP1 scored only minimally in our screens, suggesting that PIK3CA^mut^ tumors may be selectively dependent on RHEB-mTORC1 versus mTORC2, as has been recently suggested (37).

Active KRAS predominantly signals for growth and proliferation through the MAPK signaling pathway, though it can also activate the PI3K pathway (38). Comparing lethal gene pairs in KRAS^mut^ versus BRAF^V600E^ lines revealed significant differences in digenic dependencies, despite both oncogenes activating MAPK signaling (**Fig. 2F**). As before, we observed that KRAS-containing gene pairs were the most effective pairs in KRAS^mut^ lines; conversely, gene pairs including MAP2K1, MAPK1, or IRS2 were highly lethal in BRAF^V600E^ lines (**Fig. 2C, D and F**). Interestingly, MAP2K1_MAPK1 was highly effective in BRAF^V600E^ lines (LFC -3.84), but only modestly so in KRAS^mut^ lines (LFC -0.99), despite both oncogenes activating downstream MAPK signaling.

To assess this further, we conducted specific knockouts of MAP2K1 and MAPK1. Consistent with the screens, dual knockout of MAP2K1 and MAPK1 demonstrated significantly greater growth inhibition in BRAF^V600E^ versus KRAS^mut^ lines across multiple lineages and across lines not included in the screens (**Figs. 2G, S5A**). Combinatorial siRNA knockdown experiments yielded similar results, excluding artifactual effects of CRISPR knockout (**Fig. S5B**).

MEK1/2 inhibitors are active in BRAF^mut^ cancers; however, significant dose-limiting skin, gastrointestinal, and ocular toxicities limit their efficacy (39). We compared MAP2K1_MAPK1 (MEK1/ERK2) to the paralog pairs MAP2K1_MAP2K2 (MEK1/2) and MAPK1_MAPK3 (ERK1/2) in BRAF^V600E^ versus all other lines in our screens. In BRAF^V600E^ lines, MAP2K1_MAPK1 was comparable to both dual paralog knockouts (p = 0.95), whereas in all other lines, MAP2K1_MAPK1 was substantially less lethal than the dual paralog knockouts (p = 0.017) (**Fig. 2H**). This selective profile suggests that a combination therapy targeting MEK1 and ERK2 while avoiding their paralogs could increase tolerability and efficacy in BRAF^V600E^ cancers.

### Dual loss of KRAS and FGFR signaling is synergistic in KRAS^mut^ PDAC

We next asked whether digenic dependencies are influenced by tumor lineage. In the screening cohort, four of the twelve lines were KRAS^mut^ PDAC lines (HPAFII, PANC0203, PANC1, SNU410), and all four are dependent upon KRAS (40). Consistently, KRAS_AAVS1 guide pairs were highly inhibitory in these lines (mean LFC of -3.2) (**Fig. 3A**) and were more growth inhibitory than 98.5% of all gene pairs in the KRAS^mut^ PDAC lines, with only 150 gene pairs demonstrating greater lethality than KRAS_AAVS1. Nearly all (93.3%; 140/150) gene pairs with greater lethality than KRAS_AAVS1 contained KRAS.

**Figure 3.**
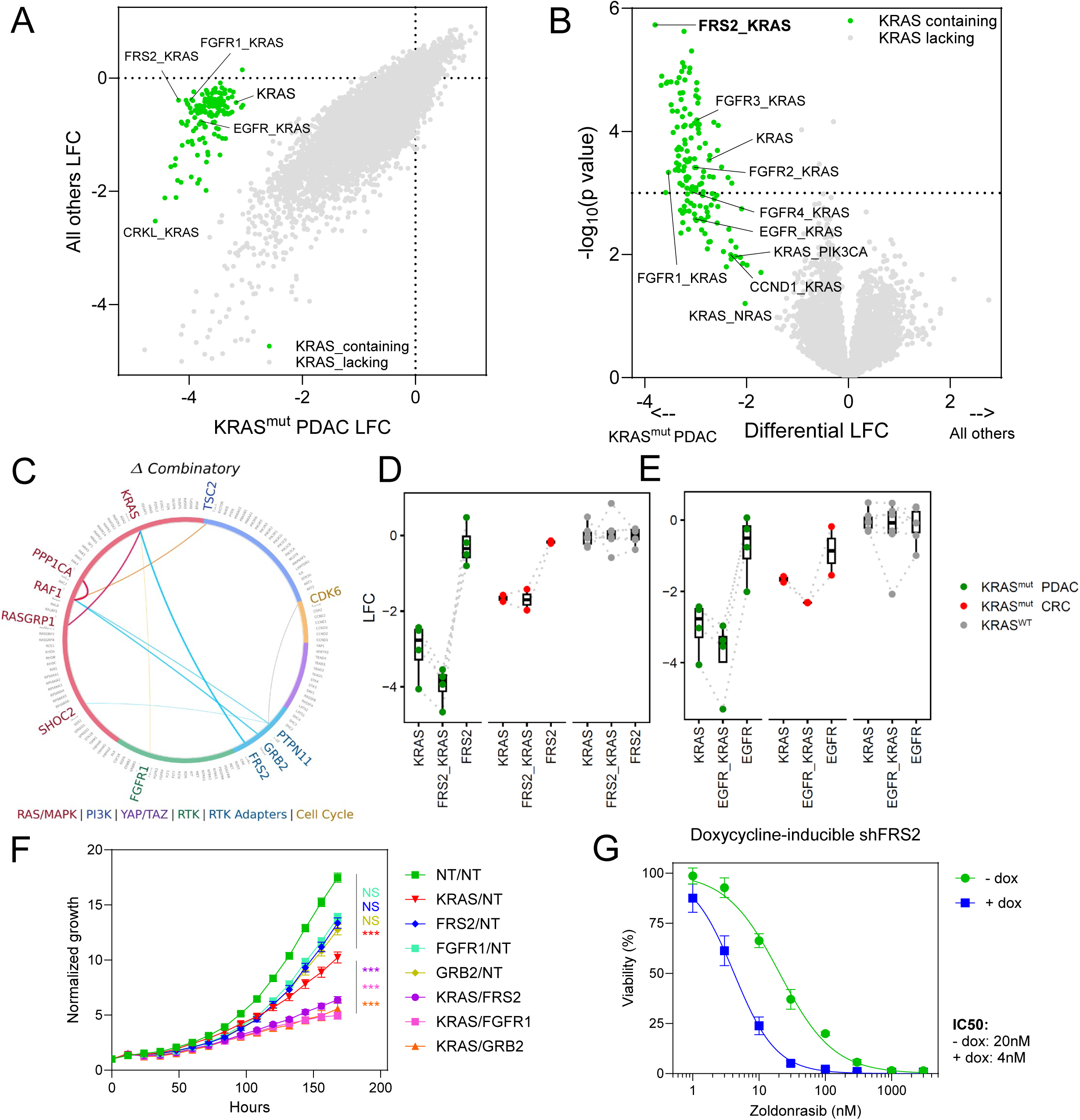
KRAS_FRS2 emerges as the top digenic dependency in KRAS^mut^ PDAC. (A) LFC plot in KRAS^mut^ PDAC versus all other lines. (B) Volcano plot of differential LFC in KRAS^mut^ PDAC versus all other lines. Y-axis shows statistical significance (-log_10_ p value). Horizontal dashed line marks p = 0.001. (C) Differential synergy map showing all gene pairs with differential Combinatory score ≤ -1 in KRAS^mut^ PDAC versus all other lines. (D-E) Boxplots display LFC scores for KRAS_FRS2 (D) and KRAS_EGFR (E) and corresponding single genes, separated into KRAS^mut^ PDAC, KRAS^mut^ CRC, and KRAS^WT^ lines. (F) siRNA dual knockdown of KRAS and FGFR-pathway genes leads to significantly reduced growth versus single knockdown in SNU410 cells. Cell growth was assayed using Incucyte and normalized to timepoint 0. Data represent mean ± SD. *P* values calculated by extra sum-of-squares *F* test comparing growth rate constants of exponential growth curve fits. Single gene knockdowns were compared versus siNT/siNT, and dual gene knockdowns were compared versus siKRAS/siNT. ***: *P* < 0.001, NS: *P* > 0.05. (G) Zoldonrasib dose-response curve of PANC0203 cells with or without doxycycline induction of an shRNA targeting FRS2. Viability measured by CellTiter-Glo. Data represent mean ± SD.

By comparing the four KRAS^mut^ PDAC screens versus all eight other screens (**Figs. 3B, S6A**), we found that KRAS gene pairs were generally more effective in KRAS^mut^ PDAC; however, 20.9% (30/143) of KRAS gene pairs, such as KRAS_NRAS, KRAS_CCND1, and KRAS_PIK3CA, were effective in the other screens and demonstrated *less* selectivity for KRAS^mut^ PDAC than knockout of KRAS alone. Gene pairs that were the most enriched in KRAS^mut^ PDAC included multiple members of the FGFR signaling pathway in combination with KRAS, with KRAS_FRS2 being the most effective and statistically significant (ΔLFC: -3.8; p = 1.86*10^-6^). FRS2 is the adaptor for all FGF-receptors (FGFR1-4) and couples FGFR signaling to RAS/MAPK activation via GRB2/SOS, leading to RAS^WT^ activation; as such, FRS2 knockout may serve as a surrogate for pan-FGFR inhibition. Among the FGFR paralogs, KRAS_FGFR1 was the most lethal, though KRAS_FGFR2, KRAS_FGFR3, and KRAS_FGFR4 were all more lethal than KRAS alone, implying that the maximal lethality of KRAS_FRS2 may be due to impairing the function of multiple FGFR paralogs. Similarly, KRAS_FRS2 and KRAS_FGFR1 were among only nine gene pairs with strong differential combination effect in KRAS^mut^ PDAC (ΔCombinatory score <-1) (**Fig. 3C**).

The individual screen results showed that KRAS_FRS2 was synergistic in all four KRAS^mut^ PDAC lines screened, including both epithelial (HPAFII, PANC0203) and mesenchymal (PANC1, SNU410) lines (**Fig. 3D**). In contrast, KRAS_FRS2 had no additional lethality over KRAS alone in KRAS^mut^ CRC, and neither individual gene nor the gene pair had significant lethality in KRAS^WT^ lines. Similar results were seen with KRAS_FGFR1, although only 3 of 4 KRAS^mut^ PDAC lines demonstrated a Combinatory score <-0.5 for this combination (**Fig. S6B**). By comparison, KRAS_EGFR was a synergistic combination in KRAS^mut^ CRC lines but had a more muted effect than that observed for KRAS_FRS2 or KRAS_FGFR1 in KRAS^mut^ PDAC lines (**Figs. 3E, S6C**). These data are consistent with the clinical role of combining KRAS^G12C^ inhibition with an EGFR-blocking antibody in KRAS^G12C^ CRC (10,11) and further support a significant, PDAC-specific role for FGFR signaling.

To validate these CRISPR-based findings, we assessed the effects of combined siRNA knockdown of KRAS and FRS2, FGFR1, or GRB2, an adapter protein that promotes FRS2-mediated activation of MAPK or PI3K signaling (41). In both PANC0203 and SNU410, KRAS knockdown substantially inhibited growth, while each of FRS2, FGFR1, or GRB2 knockdown had little to no growth inhibition (**Figs. 3F, S6D-E**). In contrast, the knockdowns of KRAS/FRS2, KRAS/FGFR1, and KRAS/GRB2 demonstrated substantially greater growth inhibition compared to KRAS. Consistently, knockdown of FRS2 or FGFR1 sensitized PANC0203 to the covalent KRAS^G12D^-selective inhibitor zoldonrasib (42,43) by more than 4-fold (**Figs. 3G, S6F-G**). Taken together, these experiments support the RAS+ screen results, identifying a novel genetic interaction between KRAS and FGFR in KRAS^mut^ PDAC.

### Combined small molecule inhibition of KRAS and FGFR is synergistic in KRAS^mut^ PDAC

To determine the therapeutic relevance of these findings, we turned to combinatorial small molecule inhibition. To minimize potential combinatorial toxicity, we chose to study the KRAS^G12D^ selective inhibitor zoldonrasib, which has demonstrated minimal toxicity and promising efficacy in KRAS^G12D^ PDAC in Phase 1 trials (43). Combinations of zoldonrasib with multi-FGFR inhibitors (FGFR1-3: pemigatinib; FGFR1-4: futibatinib, erdafitinib) and isoform-selective FGFR inhibitors (FGFR2: lirafugratinib; FGFR3: TYRA-300; FGFR4: roblitinib) demonstrated combinatorial effects with zoldonrasib in KRAS^G12D^ PDAC lines, with especially strong synergy observed in SNU410 (**Figs. 4A, S7A**). The isoform-selective FGFR inhibitors demonstrated more modest combinatorial effects. We thus selected futibatinib, which is FDA-approved for FGFR2^fusion^ cholangiocarcinoma (44), for further studies.

**Figure 4.**
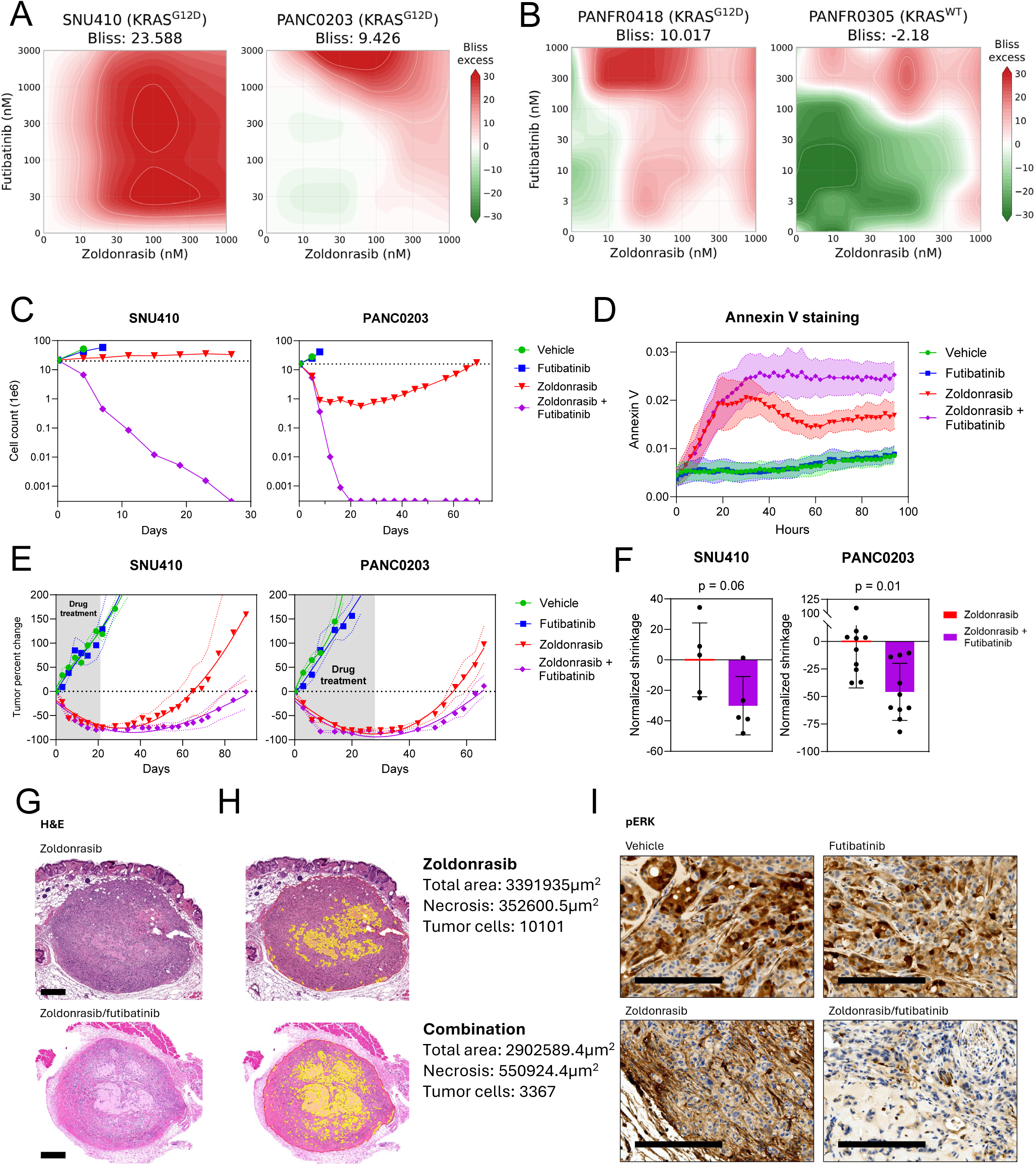
KRAS and pan-FGFR inhibition is synergistic in KRAS^mut^ PDAC. (A) Zoldonrasib x futibatinib Bliss synergy matrices for the KRAS^G12D^ PDAC 2D cell lines SNU410 (left) and PANC0203 (right). Viability was measured by CellTiter-Glo, and Bliss synergy scores were calculated across the full dose-response matrix. (B) Zoldonrasib x futibatinib Bliss synergy matrices for the PDAC organoids PANFR0418 (KRAS^G12D^; left) and PANFR0305 (KRAS^WT^; right). Viability was measured by CellTiter-Glo 3D, and Bliss synergy scores were calculated across the full dose-response matrix. (C) Large scale, longitudinal drug treatment experiments in SNU410 (left) and PANC0203 (right), treated with zoldonrasib 1000nM, futibatinib 300nM, combination, or vehicle. Cell counts are plotted over time on a log10-scaled y-axis. The dotted line indicates the starting cell number of 20*10^6^ cells; values at the bottom of the y-axis indicate no remaining cells. (D) Annexin V staining of SNU410 cells treated with zoldonrasib 1000nM, futibatinib 300nM, combination, or vehicle. Annexin V staining was assayed using Incucyte and normalized to confluence. (E) Measurements of SNU410 (left; vehicle/zoldonrasib/combination: n = 5, futibatinib: n = 3) and PANC0203 (right; all cohorts: n = 6-10) subcutaneous tumors in mice treated with zoldonrasib (100mg/kg PO QD), futibatinib (25mg/kg PO QD), combination, or vehicle. Mice were treated continuously for 3-4 weeks, followed by treatment cessation with continued tumor monitoring; the gray area on each plot represents treatment period. Dotted line indicates average tumor size for each cohort at start of treatment. (F) Tumor shrinkage of zoldonrasib versus combination treatment cohorts at 2 weeks of treatment (SNU410: left, day 15; PANC0203: right, day 14), normalized to tumor shrinkage in zoldonrasib cohorts. *P* value calculated by two-tailed unpaired Student’s *t* test. (G) Representative sections showing on-treatment PANC0203 subcutaneous tumors after staining with hematoxylin and eosin (Scale bars: 400µm). (H) Necrotic area and tumor cell count analysis on pathologist-annotated tumor bed regions in zoldonrasib- or combination-treated tumors. (I) Representative sections showing on-treatment PANC0203 subcutaneous tumors after staining with anti-phospho-ERK (brown) and counter-stained with hematoxylin (Scale bars: 200µm).

We next tested futibatinib in combination with the pan-RAS inhibitor daraxonrasib and the KRAS^G12D^ inhibitor MRTX1133, which bind different KRAS pockets. Both exhibited synergy with futibatinib, demonstrating that both mutant-selective and pan-RAS inhibitors synergize with FGFR inhibition (**Fig. S7B**). Similarly, in the KRAS^G12D^ organoid PANFR0418, the zoldonrasib/futibatinib combination was synergistic (**Fig. 4B**). In contrast, no combinatorial effect was observed in the KRAS^WT^ organoid PANFR0305. These data support the therapeutic potential of combined KRAS/FGFR inhibition.

### Combined inhibition of KRAS and FGFR is highly lethal to KRAS^mut^ PDAC cells

While the initial results demonstrated synergistic growth suppression with combined KRAS/FGFR inhibition, we next asked whether the combination could more effectively kill cancer cells than KRAS inhibition alone over longer treatment periods and in larger, more heterogeneous cell populations. To this end, we conducted long-term combination experiments starting with 2*10^7^ SNU410 or PANC0203 cells, and treating to resistance with high doses of zoldonrasib, futibatinib, combination, or DMSO control (**Fig. 4C**). In both lines, futibatinib monotherapy had minimal effects (**Fig. 4A**). In SNU410, zoldonrasib monotherapy was cytostatic but not cytotoxic, and cells continued slow, progressive proliferation throughout treatment, albeit reduced versus DMSO or futibatinib monotherapy. In contrast, the zoldonrasib/futibatinib combination was highly cytotoxic, progressively killing all cells by day 27. In PANC0203, zoldonrasib monotherapy reduced cell counts by ∼96% over two weeks; however, a small, resistant population of cells survived and grew out over two months on continuous zoldonrasib treatment. In contrast, zoldonrasib/futibatinib combination therapy rapidly killed all cells by day 20. Similarly, in a third KRAS^G12D^ PDAC line (SUIT2), zoldonrasib monotherapy eliminated ∼88% of cells, but the remaining resistant cells rapidly grew out. In contrast, the zoldonrasib/futibatinib combination eliminated nearly 99% of cells over 2 weeks, corresponding to a ∼10-fold reduction in viable cells compared with zoldonrasib monotherapy (**Fig. S8A**).

We next sought to understand the mechanisms underlying loss of cell viability. We measured Annexin V staining continuously during treatment with zoldonrasib, futibatinib, or combination. As expected, futibatinib monotherapy induced minimal Annexin V activity. In SNU410, zoldonrasib rapidly induced Annexin V activity in the first 24 hours, after which signal substantially declined over the next 24 hours and then plateaued (**Fig. 4D**). In contrast, zoldonrasib/futibatinib led to comparable Annexin V induction over the first 24 hours, which increased over the subsequent 24 hours and was sustained at a high level for at least four days. Combination therapy likewise produced greater and more sustained Annexin V signaling versus zoldonrasib monotherapy in both PANC0203 and SUIT2, matching the increased cell death seen in our long-term drug studies (**Fig. S8B-C**). The transient apoptotic response to KRAS monotherapy versus a sustained and heightened response with FGFR co-inhibition suggests a role for FGFR-mediated adaptive survival signaling in response to KRAS inhibition.

### Combined small molecule inhibition of KRAS and FGFR is synergistic *in vivo*

We next tested the zoldonrasib/futibatinib combination in SNU410 and PANC0203 subcutaneous xenografts. Mice were treated with zoldonrasib, futibatinib, combination, or vehicle at drug concentrations reported to be efficacious and tolerable as monotherapies (42,45). Treatment continued for 3-4 weeks, followed by treatment discontinuation and continued tumor monitoring. Monotherapy and combination treatments were well-tolerated, with all cohorts maintaining comparable weight during and after treatment (**Fig. S8D-E**). As in the *in vitro* data, futibatinib had no effect on tumor growth, and both futibatinib monotherapy and vehicle-treated tumors grew through treatment (**Fig. 4E**). In contrast, zoldonrasib monotherapy led to robust tumor regression. However, after treatment cessation, zoldonrasib-treated tumors regrew, with tumors reaching starting size after 44 days (SNU410) and 28 days (PANC0203) off-treatment. The addition of futibatinib to zoldonrasib led to marked tumor regression, achieving more rapid and deeper tumor regressions as compared to zoldonrasib monotherapy after two weeks on treatment (**Fig. 4F**). Additionally, once treatment was held, tumor regrowth was suppressed for significantly longer with combination therapy with tumor regrowth delayed by 57% (69 total days) in SNU410 and 36% (38 total days) in PANC0203 compared to zoldonrasib monotherapy (**Fig. 4E**). Thus, time-limited treatment with combination zoldonrasib/futibatinib deepened responses and delayed tumor outgrowth versus zoldonrasib monotherapy without evidence of systemic toxicity.

To assess proximal treatment-related effects, we collected tumor samples after two weeks, while zoldonrasib- and combination-treated tumors were still responding. We observed modest treatment effect with limited acellular debris in the zoldonrasib-treated tumors (**Fig. 4G**, top panel). In contrast, combination-treated tumors had prominent acellular debris (**Fig. 4G**, bottom panel), suggesting that caliper measurements might underestimate combination efficacy. Indeed, quantitative image analysis of pathologist-annotated tumor beds demonstrated a greater necrotic fraction in combination-treated tumors, with roughly 3-fold reduction in tumor nuclei despite comparable tumor bed areas (**Fig. 4H**). Pharmacodynamic immunohistochemistry measures showed downregulation of pERK in zoldonrasib- and combination-treated tumors (**Fig. 4I**). Gene set enrichment analysis (GSEA) on RNA-seq data from on-treatment tumors demonstrated downregulation of growth and proliferation pathways in all three treatment arms relative to vehicle which was most pronounced in zoldonrasib- and combination-treated tumors (**Fig. S8F; Table S5**). Zoldonrasib-treated tumors also showed upregulation of type I and II interferon response gene sets, expression of which has previously been associated with generalized therapy resistance (46,47); this zoldonrasib-induced interferon response was lost upon the addition of futibatinib, suggesting the elimination of an interferon-high, FGFR-dependent resistant population. Zoldonrasib- and combination-treated tumors also showed upregulation of FGFR1 and FGFR2 transcripts, suggesting that these FGFR paralogs may confer resistance (**Fig. S8G**). This was especially notable for FGFR2, which was undetectable in vehicle- and futibatinib-treated tumors, but substantially elevated in both zoldonrasib- and combination-treated tumors.

### KRAS inhibition induces FGFR activation

Having observed synergy between KRAS and FGFR inhibition *in vitro* and *in vivo*, we sought to understand the mechanisms underlying this gene interaction. To this end, we treated PANC0203 and SNU410 cells with extended durations of suppressive doses of zoldonrasib. As previously described, KRAS inhibition initially suppressed MAPK signaling (pERK) (42); however, we observed recovery of pERK signaling by day 10 in PANC0203 and day 3 in SNU410 (**Fig. 5A**). This was accompanied by concurrent upregulation of pFGFR and PI3K signaling (pAKT), suggesting that activation of FGFR signaling might reactivate proliferative signaling in the setting of KRAS inhibition.

**Figure 5.**
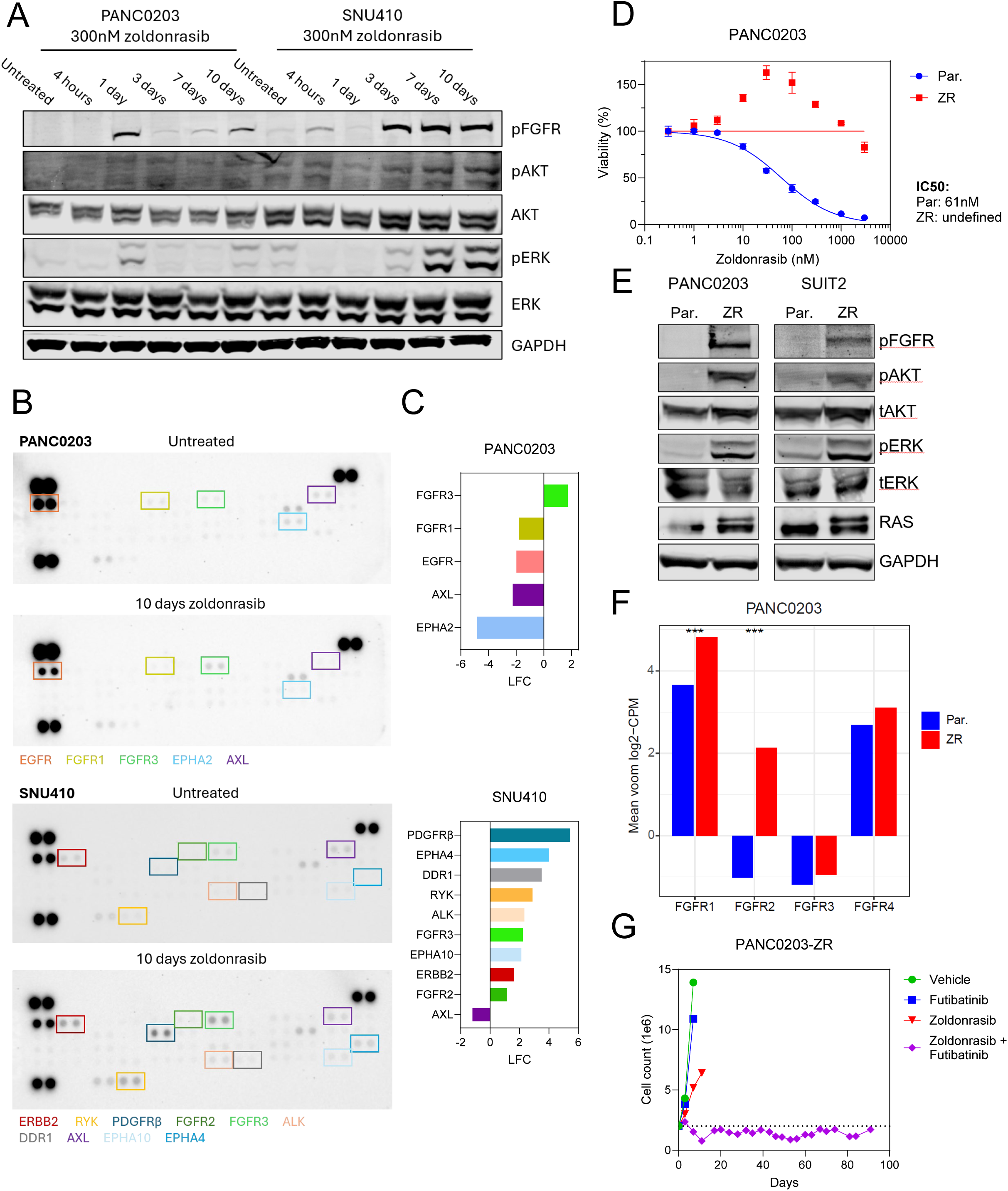
FGFR activation causes KRAS inhibitor resistance and contributes to acquired resistance. (A) Immunoblot for pFGFR, pERK, and pAKT of PANC0203 and SNU410 cells treated with zoldonrasib 300nM for increasing durations. (B) Phospho-RTK arrays of PANC0203 and SNU410 cells treated with zoldonrasib 300nM for 0 or 10 days. pRTKs with at least two-fold increase or decrease in signal intensity are highlighted. (C) pRTKs with at least two-fold increase or decrease in signal intensity from (B). (D) Zoldonrasib dose-response curve of PANC0203 parental versus zoldonrasib-resistant (ZR) cells. Viability measured by CellTiter-Glo. Data represent mean ± SD. (E) Immunoblots for pFGFR, pERK, and pAKT of PANC0203 (left) and SUIT2 (right) parental versus ZR cells. (F) Bar plot showing mean voom-normalized log2 CPM for FGFR1-4 in PANC0203 parental and ZR cells. Data represent mean expression across biological replicates. *P* values calculated using empirical Bayes moderated *t* test. ***: *P* < 0.001. (G) Large scale, longitudinal drug treatment experiments in PANC0203-ZR, treated with zoldonrasib 1000nM, futibatinib 300nM, combination, or vehicle. The dotted line indicates the starting cell number of 2*10^6^ cells.

To evaluate RTK activation after prolonged KRAS inhibition, cell lysates extracted from PANC0203 and SNU410 cells before and after 10 days of zoldonrasib treatment were hybridized to antibody arrays detecting phosphorylation of 49 RTKs to assess RTK phosphorylation (**Fig. 5B; Table S6**). In PANC0203, which underwent substantial cell death over 10 days, only FGFR3 showed increased phosphorylation, suggesting a key role for FGFR signaling in the surviving cells. In SNU410, which showed slower but continuous growth despite KRAS inhibition, we observed increased phosphorylation of 9 RTKS, including both FGFR2 and FGFR3 (**Fig. 5C**).

### Upregulated FGFR signaling in isogenic KRAS inhibitor resistant models is required for maintaining inhibitor resistance

To characterize mechanisms contributing to zoldonrasib resistance, we generated PANC0203 and SUIT2 populations with acquired resistance to high-dose zoldonrasib (as described in **Figs. 4C** and **S8A**). We termed these zoldonrasib-resistant (ZR) lines and maintained them in zoldonrasib. ZR lines had zoldonrasib IC50s >3µM, representing >10-fold resistance (**Figs. 5D, S9A**). Both lines also demonstrated cross-resistance to MRTX1133 and daraxonrasib, confirming that resistance was RAS-mediated and not binding pocket-dependent (**Fig. S9B-E**). Mechanistically, immunoblots demonstrated strong upregulation of pFGFR in both ZR lines, along with corresponding upregulation of pERK and pAKT, suggesting that FGFR signaling was activated in ZR lines and contributing to proliferation and survival (**Fig. 5E**). Interestingly, higher levels of pERK were observed in each ZR line compared to the untreated parental cells, despite ongoing zoldonrasib treatment, suggesting “hyperactivation” of RAS/MAPK signaling in the ZR lines. This hyperactivation appeared to lead to RAS/MAPK-mediated oncogene toxicity, as both ZR lines showed improved growth in the presence of low doses of zoldonrasib versus in the absence of zoldonrasib (**Figs. 5D, S9A**) (48). RNA-seq data revealed that both ZR lines had strong upregulation of FGFR1 and FGFR2 compared to the parental lines, consistent with our *in vivo* data demonstrating upregulation of FGFR1/FGFR2 in zoldonrasib-treated tumors (**Figs. 5F, S9F**). Finally, we treated large populations (2*10^7^) of ZR cells with zoldonrasib, futibatinib, combination, or DMSO. As expected, ZR cells grew despite zoldonrasib treatment, and futibatinib had minimal impact on cell growth. In contrast, combination therapy prevented ZR cell outgrowth for >3 months (**Fig. 5G**). These data suggest that FGFR activation mediates the RAS inhibitor resistance phenotype and show that combined KRAS/FGFR inhibition can overcome or blunt acquired resistance to KRAS inhibitors.

### Cancer-associated fibroblasts are a primary source of FGF ligands in PDAC tumors

Each of the four FGFR paralogs requires specific FGF ligands for its activation, and there are 18 secreted FGF ligands, each having a different FGFR activation profile (49). The finding that FRS2 was the top combination partner with KRAS suggested that FGFR activation in PDAC might be mediated by FGF ligands signaling through multiple FGFRs and converging on FRS2. As such, we sought to identify which FGF ligands are present in PDAC tumors and which cells in the tumor or TME produce those FGF ligands. Prior work has suggested that FGF ligand production can be either autocrine from PDAC cells or paracrine from CAFs, which make up a large portion of the PDAC TME (50–52). To assess this, we conducted a meta-analysis of five large patient single-cell RNA-sequencing (scRNA-seq) studies that included both PDAC and TME cells. In total, we analyzed 364,082 cells from 97 patients (53–57). We visualized the unified dataset via UMAP, recovering the PDAC cellular ecosystem with resolution of malignant, fibroblast, vascular and immune populations (**Fig. 6A; Methods**). Cells from the five cohorts were intermixed, indicating appropriate data integration (**Fig. S10A**), and non-malignant compartments intermixed across the dataset, while the malignant compartment showed patient-specific clustering, as expected (**Fig. S10B**). Projecting individual FGF ligands onto this embedding revealed CAF-specific, high expression of FGF1, FGF7, and FGF10, as well as expression of FGF2 in both the PDAC and CAF compartments (**Fig. 6B**); interestingly, several of these ligands (FGF7, FGF10) are mesenchyme-derived factors which signal to epithelial receptors driving differentiation and outgrowth in early pancreatic embryogenesis, suggesting a rationale for their continued or re-activated mesenchymal expression in PDAC (58).

**Figure 6.**
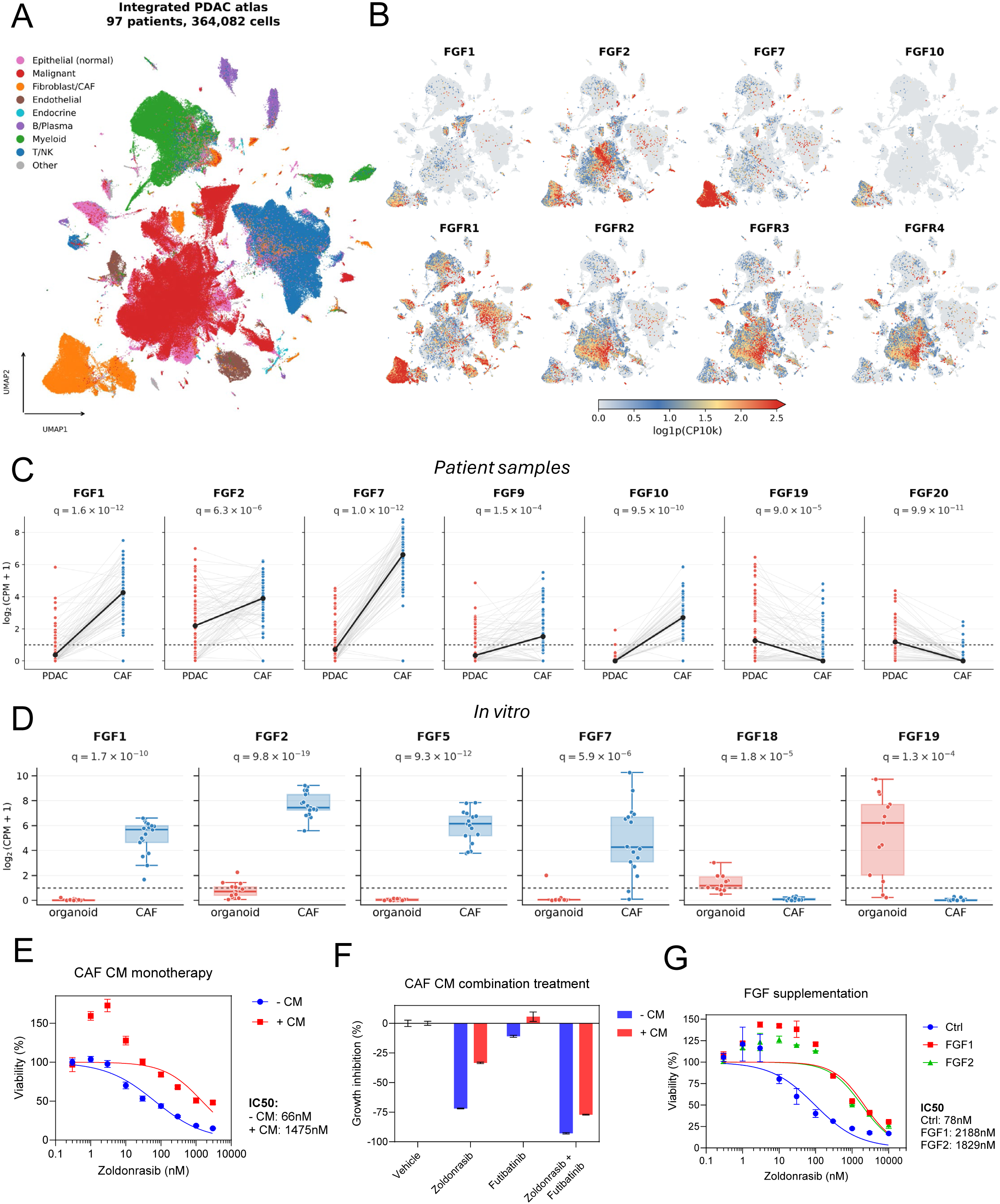
The CAF compartment is the primary source of FGF ligand in PDAC tumors. (A) UMAP of 364,082 cells from 97 patients across five PDAC scRNA-seq cohorts (53–57), colored by cell type. Harmony batch integration was performed across cohorts. (B) Normalized expression of select FGF ligands and FGFRs overlaid on the UMAP embedding shown in A. (C) Patient-level comparison of FGF ligand pseudobulk expression between PDAC and CAF compartments for all secreted FGF ligands with significantly different expression. Bolded black lines connect compartment medians. *Q* values calculated via paired Wilcoxon signed-rank test across patients followed by Benjamini-Hochberg correction. Dotted line represents CPM = 1. (D) Expression of all secreted FGF ligands with significantly different expression in 17 CAF versus 13 PDAC organoid models. *Q* values calculated via Welch’s two-sample t-test followed by Benjamini-Hochberg correction. Dotted line represents CPM = 1. (E) Zoldonrasib dose-response curve of SNU410 supplemented with or without CAF conditioned media (CM). Viability measured by CellTiter-Glo. Data represent mean ± SD. (F) Growth inhibition of zoldonrasib 300nM, futibatinib 300nM, combination, or vehicle in SNU410 supplemented with or without CAF CM. Viability measured by CellTiter-Glo. Data represent mean ± SD. (G) Zoldonrasib dose-response curve of SNU410 with supplementation of FGF1, FGF2, or vehicle. Viability measured by CellTiter-Glo. Data represent mean ± SD.

We further explored FGF levels by quantifying mRNA expression of each secreted FGF ligand in the malignant and CAF compartments. Several FGF ligands, including FGF1/FGF2/FGF7/FGF9/FGF10, demonstrated higher bulk expression in CAFs than PDAC cells (**Fig. S10C**). Notably, this pattern was maintained in both primary and metastatic tumors, highlighting that CAFs may be the primary source of FGF ligands even after PDAC cells have disseminated from the pancreas. To formally compare expression, we constructed per-patient pseudobulk profiles of the CAF and malignant compartments. Here, several FGF ligands had significantly greater expression in CAFs: FGF1 (ΔLFC 3.9; q = 1.6*10^-12^), FGF2 (ΔLFC 1.7; q = 6.3*10^-6^), FGF7 (ΔLFC 5.9; q = 1.0*10^-12^), FGF9 (ΔLFC 1.2; q = 1.5*10^-4^), and FGF10 (ΔLFC 2.5; q = 9.5*10^-10^) (**Fig. 6C**). In contrast, the only FGF ligands more highly expressed in malignant cells were FGF19 (ΔLFC -1.3; q = 9.0*10^-5^) and FGF20 (ΔLFC -1.2; q = 9.9*10^-11^), and both had relatively low median malignant expression.

This meta-analysis also allowed the examination of FGFR expression across the integrated dataset. Within the malignant compartment, all FGFR paralogs were expressed, with FGFR1 having the lowest aggregate expression (**Figs. 6B, S10C**). In contrast, FGFR1 was predominant in CAFs, which had substantially less FGFR2 and minimal FGFR3/4 expression. We also assessed the extent to which multiple FGFRs were expressed in any individual patient’s tumor, finding that most patients have detectable tumor levels of all or nearly all FGFRs (**Fig. S10D**).

To determine whether in vitro models reflect these patient data, we leveraged a panel of 17 primary CAF lines that were isolated from patient tumors, as well as a panel of 13 PDAC organoids, isolated from 7 patient and 6 PDX tumors (Yeh et al, manuscript in preparation) (59,60). We collected RNA-seq data across the panel, finding that, similar to the patient data, FGF1, FGF2, FGF5, and FGF7 were more highly expressed in CAFs, while only FGF18 and FGF19 were more highly expressed in organoids (**Figs. 6D, S11A; Table S7**). Among four paired CAF and organoid lines derived from the same tumors, FGF1, FGF2, and FGF5 were elevated in CAFs, while FGF18 and FGF19 were elevated in organoids (**Fig. S11B**). Also matching our patient analysis, FGFR1 was more highly expressed in CAFs (though still expressed in most organoids), while FGFR2-4 were more highly expressed in organoids (**Fig. S11C**). The concordance between the patient samples and the in vitro models suggests that immortalized CAFs retain the FGF/FGFR signaling axis observed in patient tumors.

### PDAC CAFs promote KRAS inhibitor resistance in malignant cells

We next assessed whether CAFs can contribute to KRAS inhibitor resistance via paracrine signaling. To do so, we assessed drug sensitivity in PDAC cells exposed to conditioned media (CM) from an immortalized PDAC CAF line. CM induced substantial resistance, with up to 22x shifts in zoldonrasib IC50s (**Figs. 6E, S12A**). As in the ZR lines, cells exposed to CM showed growth advantage at low doses of zoldonrasib versus no zoldonrasib, consistent with CM-mediated RAS/MAPK hyperactivation. Immunoblotting demonstrated that CM exposure increased pFGFR, with associated increases in pERK and/or pAKT (**Fig. S12B**). Importantly, zoldonrasib/futibatinib overcame CM-induced resistance, with the combination more than doubling the effect of zoldonrasib alone in CM-treated cells. These data suggest that FGF-mediated signaling is causal for the observed KRAS inhibitor resistance induced by CM (**Figs. 6F, S12C**). Notably, while futibatinib overcame the majority of CM-mediated resistance, the combination was modestly less effective in CM-exposed versus CM-naïve cells, suggesting that other CAF-mediated factors may also contribute to KRAS inhibitor resistance, as has been reported (61).

### FGFR activation causes KRAS inhibitor resistance

Finally, we sought to understand whether individual FGFs were sufficient to induce KRAS inhibitor resistance, and we exogenously supplemented FGF1 or FGF2 in cells treated with zoldonrasib. FGF1 or FGF2-treated cells demonstrated significant resistance to KRAS inhibition, with up to 20x shifts in zoldonrasib IC50s (**Figs. 6G, S13A**). Similarly, lentiviral overexpression of FGF1 or FGF2 strongly induced zoldonrasib resistance, with comparable IC50 shifts to exogenous supplementation (**Fig. S13B-C**). Consistently, FGFR inhibition with futibatinib fully overcame the resistance induced by FGF overexpression, with the zoldonrasib/futibatinib combination demonstrating extremely high levels of synergy in FGF2-overexpressing cells (**Fig. S13D**).

## DISCUSSION

Targeted therapy has reshaped the therapeutic landscape of many cancers, but adaptive signaling and acquired resistance remain barriers to durable benefit. In this study, we developed the RAS+ combinatorial CRISPR knockout library to map genetic interactions across key oncogenic signaling pathways that commonly mediate targeted therapy resistance.

A central finding from our screens is that genetic and lineage context heavily shapes genetic interaction distributions. KRAS^mut^, BRAF^mut^, and PIK3CA^mut^ cancers each displayed distinct lethality maps despite substantial overlap in oncogenic signaling pathways. Importantly, in KRAS^mut^ models, the most effective combinations were overwhelmingly centered on KRAS itself; this suggests that efficacious clinical combinations will require direct suppression of KRAS paired with blockade of resistance nodes rather than combinations lacking KRAS inhibition. It further implies that resistance to KRAS inhibition in KRAS^mut^ tumors might be more common at the level of KRAS than in downstream pathway components, as has been recently reported (18,62). In sum, these results establish that digenic dependencies can reveal context-specific signaling principles that cannot always be inferred from linear signaling hierarchy.

Emblematic of this is the observation that MEK1/ERK2 co-suppression is selectively lethal in BRAF^V600E^ cells. Unfortunately, no paralog-selective MEK or ERK inhibitors exist yet due to the similarity of binding pockets in MEK1/MEK2 and in ERK1/ERK2. However, targeted protein degradation may offer a route to paralog selectivity, as degrader specificity is governed by ternary complex geometry rather than strictly by domain binding affinity. For example, the CDK4/6 inhibitor palbociclib inhibits both CDK4 and CDK6, but a palbociclib-derived heterobifunctional degrader selectively degrades CDK6 while sparing CDK4 (63). This principle is being explored for ERK, and several ERK1/2 inhibitors act as ERK2 but not ERK1 degraders (64); our findings provide rationale for continued pursuit of paralog-specific MEK1 and ERK2 degraders which may yield a wider therapeutic index than dual-paralog inhibition.

More broadly, we identified recurrent, highly lethal combinations between core MAPK nodes and proximal RTK adapters, including IRS2 paired with BRAF, MAP2K1, or MAPK1 in BRAF^mut^ lines and FRS2 paired with KRAS in KRAS^mut^ PDAC. These findings suggest a model in which disruption of oncogenic signaling exposes dependence on upstream RTK inputs converging through adapter proteins, with relevant input(s)/adapter(s) shaped by genotype, lineage, or feedback circuitry. Consistent with this model, insulin/IGF receptor-PI3K signaling has been implicated in adaptation and resistance to BRAF inhibition in melanoma and CRC (65–67).

The identification of FRS2, a convergent adapter downstream of FGFR1-4, as the most selective KRAS combination partner in KRAS^mut^ PDAC was unexpected but may reflect re-engagement of lineage-encoded developmental programs. During pancreatic organogenesis, mesodermal FGF2 acts on epithelial FGFRs to permit expression of the master pancreas transcription factor PDX1 (68), after which mesenchymal FGF7/10 act on epithelial FGFR2b to expand PDX1+ progenitors into pancreatic tissue (58,69). FGF-FGFR signaling therefore constitutes a canonical program in the developing pancreas. This circuitry remains accessible in neoplastic pancreatic cells: FGF10 sustains propagation of malignant pancreatic organoids (70,71), while FGFR2 is progressively upregulated during KRAS-driven pancreatic metaplasia and transformation (52).

Our data suggest that this existing FGF-FGFR circuitry can be coopted to drive resistance. Through a comprehensive analysis of patient samples and recently derived models, we find that CAFs are likely the dominant source of FGF ligands in PDAC, and three of the most differentially expressed ligands in patient CAFs (FGF2/7/10) are those relevant for early pancreatic development. Indeed, we find that individual FGF ligands produced by CAFs are sufficient to induce KRAS inhibitor resistance and that PDAC cells themselves respond to KRAS inhibition with significant, sustained upregulation and activation of FGFR1/2. Thus, stromal ligand supply and tumor cell adaptation converge to reinforce FGFR signaling, blunting response to KRAS inhibition; FGFR co-inhibition overcomes this adaptation, deepening and prolonging tumor responses. Taken together, these findings support a model in which PDAC CAFs do not just act as generic stroma but partially reconstitute a mesenchymal niche resembling that which supports embryonic pancreatic development; PDAC cells may then exploit this niche to survive KRAS inhibition. While our analyses suggest that that the majority of FGF ligand is CAF-derived, the dependence on FGFR signaling seen in our CRISPR screens suggest that autocrine production of FGF ligands by tumor cells may also contribute to KRAS inhibitor resistance, and the absolute levels of secreted FGF by each cellular compartment remain to be determined.

Our studies demonstrated marked synergy from KRAS and pan-FGFR inhibitor combinations. Importantly, combination therapy was highly and selectively cytotoxic, driving far greater cell killing in vitro and producing deeper and more durable tumor regressions in vivo. In these studies, we anchored therapy on the mutant-selective KRAS^G12D^ inhibitor zoldonrasib rather than the pan-RAS inhibitor daraxonrasib to improve combination tolerability; indeed, the zoldonrasib/futibatinib combination was well-tolerated in murine studies. Recently, Lin and colleagues assessed RTK-driven adaptive signaling following KRAS inhibition in PDAC, reporting FGFR signaling as a mediator in mesenchymal models (72), consistent with prior work identifying FGFR1-mediated adaptive resistance to MEK inhibition in KRAS^mut^ NSCLC (73) and linking this mechanism to mesenchymal state (74). These studies provide independent support for the FGFR axis identified here, and our data expand upon these findings to show that FGFR-pathway adaptation is not limited to mesenchymal disease: KRAS_FRS2 emerged as a top hit in both epithelial and mesenchymal PDAC RAS+ screens, and adaptive FGFR activation led to acquired resistance in epithelial PANC0203 and SUIT2 cells. We thus propose a model in which FGFR-mediated adaptation acts as a KRAS inhibitor resistance mechanism across PDAC cell states.

These results have several implications for clinical development. First, combined KRAS/FGFR inhibition may warrant evaluation as a strategy to deepen responses and delay resistance in KRAS^mut^ PDAC, especially using mutant-selective KRAS inhibitors to improve combination tolerability. Second, given the expression of multiple FGFRs in most patient tumors, multi-FGFR inhibitors may be necessary to overcome redundancy in adaptive signaling. Third, biomarker development should therefore assess both stromal FGF ligand expression and tumor FGFR activation, ideally through spatial or compartment-resolved analysis of paired pretreatment and on-/post-treatment biopsies.

There are several limitations to this work. Because assessing all pairwise combinations leads to quadratic library size increases, the RAS+ library was limited to 144 genes in core oncogenic signaling pathways; thus, genetic interactions involving pathways or signaling programs outside of this set (e.g., WNT, apoptosis, etc.) have not been assessed. The library also evaluates pairwise but not higher-order combinations, and deep, durable clinical regimens may require suppression of three or more nodes. Our screening cohort consisted of twelve gastrointestinal and skin lines, and our genotype- and lineage-defined comparisons thus rest on modest numbers, requiring confirmation in larger, pan-cancer screening efforts. Finally, our studies employed subcutaneous xenografts in immunodeficient mice, in which the stromal compartment is not pancreas-derived and adaptive immunity is absent; evaluation in orthotopic, immunocompetent models will be needed to understand the full contributions of the pancreatic TME.

In summary, this work establishes the RAS+ screening system to identify therapeutically relevant genetic interactions and identifies FGFR signaling as a mediator of KRAS inhibitor resistance in PDAC, in part mediated by lineage-specific aspects of the pancreatic TME. By integrating combinatorial functional genomics, pharmacologic validation, and resistance modeling, we nominate combined KRAS/FGFR inhibition as a rational and novel therapeutic strategy for KRAS^mut^ PDAC.

## METHODS

### RAS+ gene and gene pair selection

The RAS+ library was designed to target genes in the RAS/MAPK, PI3K, and YAP/TAZ pathways, as well as upstream RTKs and RTK adapters and downstream cell cycle genes. Genes were curated from the literature, including RAS pathway gene maps developed by the NIH RAS Initiative. This yielded a list of 144 genes. All pairwise gene pairs from these 144 genes were included in the RAS+ library, yielding 10,296 total gene pairs.

Each gene is targeted by 4 sgRNAs, and each gene pair is targeted by 8 sgRNA pairs. The library includes positive and negative controls: 145 pan-essential genes and 143 non-essential genes from the DepMap Avana screens dataset.

### RAS+ sgRNA design and selection

sgRNAs for the RAS+ library were curated from two well-validated CRISPR libraries—our previously generated PARADIGM library and the Broad Institute Avana library—and supplemented by de novo design for genes lacking enough qualifying guides. To identify guides with strong on-target activity, we trained a random forest classifier on the Avana 23Q4 dataset to discriminate sgRNA pairs targeting the same gene from pairs targeting different genes, using fourteen agreement metrics computed from their LFC profiles across cell lines. We applied this classifier to all same-gene pairs in both PARADIGM and Avana, and we used the predicted same-gene probability as the pairwise agreement score. Because of the optimizations in guide performance, we selected 4 sgRNAs per gene and 8 sgRNA pairs per gene pair for the RAS+ library; this design reduces the redundancy from the PARADIGM library, which contains 6 sgRNAs for each individual gene and 18 different sgRNA pairs for each gene pair.

We first drew sgRNAs from the PARADIGM library. We retained single-gene pairs with agreement > 0.7, picked the highest-agreement pair per gene, and discarded pairs in which both guides scored below 0.6 on Chronos-inferred efficacy. When the PARADIGM library did not yield four sufficient guides for a given gene, we drew additional guides from Avana under the same criteria. For genes that did not reach four guides across the two libraries, we designed additional sgRNAs de novo with crisprDesign (25). For these genes, we identified candidate SpCas9 spacers within coding-sequence regions of hg38, filtered to 30-80% GC content, no polyT runs, and no BsmBI or BbsI sites, and ranked them by a combined DeepHF/DeepSpCas9/RuleSet3 on-target score with iterative Bowtie off-target alignment allowing up to two mismatches. We merged all selected sgRNAs from the three sources as the final sgRNAs for the RAS+ library. The list of Cas9 sgRNA sequences, source, and pairs are listed in **Table S2**.

### Library cloning and plasmid production

As described in our previous work (16), sgRNA oligonucleotide pools were synthesized (Twist Bioscience) with BsmBI sites and appropriate overhang sequences spanning the two 20-nt crRNAs. Two BbsI sites were designed between the two crRNAs. The final oligonucleotide sequence was thus: 5′-AGGCACTTGCTCGTACGACGCGTCTCGCACCG [crRNA, 20nt] GTTTCAGTCTTCCGGCGAAGACACCTGAAAC [reverse complement crRNA, 20nt] CGGGAAGAGACGTTAAGGTGCCGGGCCCACAT-3′. The oligonucleotide pools were amplified with 24 cycles of PCR, purified by spin-column, digested with Esp3I, and ligated into Esp3I-digested pWRS_1001 (Addgene, Catalog No. 192205) vector via 100 cycles of Golden Gate assembly. For the incorporation of the tracrRNAs, the BbsI-digested tracrRNA fragment was cloned in between the dual crRNAs with a second round of Golden Gate cloning. Assembled library was electroporated into Stbl4 electrocompetent cells and expanded, and plasmid DNA (pDNA) was isolated.

Library lentivirus was produced using Lenti-X 293T cells (Takara Bio, Catalog No. 632180). Lenti-X 293T were co-transfected with sgRNA library and packaging plasmids psPAX2 (Addgene, Catalog No. 12260) and pMD2.G (Addgene Catalog No. 12259) using LT-1 transfection reagent (Mirus, Catalog No. MIR2300). Viral supernatants were collected 48 hours post-transfection, filtered through 0.45 μm filters, and stored at -80 °C. All lentiviral work was performed under BSL-2+ conditions with institutional biosafety approval.

### RAS+ library screening, DNA extraction and amplification, and sequencing

Screening followed protocols described in our previous work (16), with library-specific modifications. Cas9-expressing stable cell lines were generated via transduction with the pLX311-Cas9 lentiviral vector (Addgene, Catalog No. 96924) or LentiV_Cas9_Blast (Addgene, Catalog No. 125592), which carries SpCas9 under the control of the EF1a promoter and confers blasticidin resistance. Prior to screening, Cas9-positive populations were enriched through blasticidin selection. Cells were then infected with the RAS+ library virus at a low multiplicity of infection ensuring ≥250x representation (MOI 0.3–0.5; approximately 70*10^6^ cells). For viral transduction, 3*10⁶ cells per well were seeded into 12-well plates in the presence of 4 μg/mL polybrene and virus. Plates were centrifuged at 2,000 rpm for 2 hours and subsequently incubated for 24 hours. After infection, cells were expanded and subjected to puromycin selection (2 μg/mL, Gibco, Catalog No. A1113803) for 3 days. Selected cells were then grown over at least 16 days to achieve at least 6 doublings, with passaging maintaining library coverage and with cell counts recorded at each passage to monitor growth dynamics. At endpoint, cells were collected by centrifugation, and pellets were stored in -80°C. Genomic DNA was extracted using the NucleoSpin Blood XL kit (Takara Bio, Catalog No. 740950) following the manufacturer’s protocol. PCR amplification of gDNA was performed in 100-μL reactions, each containing up to 10 μg of gDNA. Amplification and barcoding were carried out using P5/P7 primers (Integrated DNA Technologies) and Titanium Taq DNA polymerase (Takara Bio, Catalog No. 639242), following the manufacturer’s protocol. Each 100-μL reaction contained 50 μL of gDNA in water, 40 μL of PCR master mix, and 10 μL of a 5 μM stock of uniquely barcoded P7 primer. PCR products were purified using Agencourt AMPure XP SPRI beads (Beckman Coulter, Catalog No. A63880) according to the manufacturer’s instructions. Libraries were sequenced on an Illumina NovaSeq platform using 50 bp paired-end reads. sgRNA sequences (20–21 nt) were mapped to a reference file of all library-encoded sgRNAs to generate count data.

### RAS+ screen data processing

Demultiplexed paired-end FASTQ files were processed using PoolQ (v3.13.2). For each read pair, sgRNA sequences were extracted from the corresponding R1 and R2 reads and combined into a construct barcode, which was matched against the RAS+ library reference. Only read pairs matching a valid dual-sgRNA construct were retained for quantification. Counts were summed across sequencing lanes and aggregated by sample to generate the final construct-level read count matrix. Finally, counts from all sequencing lanes were combined for each barcode to generate the final sgRNA read count vectors representing each sample.

### Quality control

Library representation was assessed by computing the cumulative read distribution of pDNA samples, with sgRNA constructs ranked by read abundance.

To evaluate screen quality, we computed the null-normalized median difference (NNMD) between essential and non-essential genes for each screen individually. This metric was based on gene-level naive LFCs, as described in the next section. Control gene sets for single-gene perturbations were the 145 pan-essential genes and 143 non-essential genes in the library. Screens were excluded from downstream analysis if any sequenced samples in a screen had an NNMD > -2.

### Data analysis

To calculate LFCs for single and gene pair knockouts, raw sgRNA counts were first log₂-transformed with a pseudocount of 1. The pDNA reference sample was centered by subtracting its median yielding a standardized baseline. Each sequenced sample was then adjusted by aligning the median abundance of a predefined set of negative control sgRNAs to that of the pDNA reference. This procedure yielded normalized read counts for each sgRNA in both the pDNA reference and experimental replicates. LFCs were then computed by subtracting the normalized counts in the pDNA reference from each sequenced sample. LFCs for each perturbation (either a single gene or a gene pair) were calculated by taking the median of the LFCs of all sgRNAs targeting that perturbation. For screens with multiple technical replicates, the final LFC for each perturbation was obtained by taking the median across replicates.

To calculate Combinatory scores, we compare a gene pair LFC to the more lethal (i.e., lower) of the two corresponding single gene LFCs using an empirical statistical framework. Specifically, we generated a null distribution for each screen using a set of non-synergistic gene pairs, defined as pairs in which only one or neither gene is expressed in the corresponding cell line. For each screen, null distributions were constructed by permuting the Combinatory scores of these non-synergistic pairs 100,000 times. A *p*-value was then assigned to each observed gene pair based on the left-tail probability of its Combinatory score under the corresponding null distribution. False discovery rates (FDRs) were subsequently estimated with Benjamini-Hochberg correction.

To generate clustered genetic interaction maps, genes were ordered by unsupervised hierarchical clustering of their interaction profiles. Each gene was represented by its vector of Combinatory scores across all partner genes in the library. Pairwise gene-gene dissimilarity was computed using correlation distance, so genes were grouped according to similarity in the shape of their interaction profiles rather than absolute score magnitude. Hierarchical clustering was then performed with average linkage, and optimal leaf ordering was applied.

### Generation of constructs or expression vectors for target validation

#### MAP2K1 and MAPK1 knockout

sgRNAs targeting MAP2K1/MAPK1 (ACATCCTAGTCAACTCCCGT/ GCTGACCTTGAGATCACAGG) or AAVS1/AAVS1 (TCGATCCGCCCCGTCGTTCC/CAGTTGAAGCGGCTCCAATT) were cloned into pWRS_1001 as described in the Library Cloning section. Inserts were verified by Sanger sequencing. Lentivirus was produced as described in the Library Cloning section. Cells expressing Cas9 were infected with the lentivirus, in the presence of 4ug/mL polybrene. After infection, cells were selected with puromycin for 3 days. Selected cells were seeded into 96-well plates. After 5-7 days, cell viability was determined via CellTiter-Glo (Promega, Catalog No. G7571) following manufacturer’s instructions, normalizing MAP2K1_MAPK1 viability to AAVS1_AAVS1 viability for each cell line.

#### KRAS, FRS2, FGFR1, GRB2, MAP2K1, and MAPK1 siRNA knockdown

ON-TARGETplus 2.0 siRNA pools of 4 siRNAs targeting KRAS, FRS2, FGFR1, GRB2, MAP2K1, or MAPK1, or non-targeting (siNT) pools were acquired from Dharmacon. siRNA was transfected into recipient cells using DharmaFECT 2 Transfection Reagent per the manufacturer’s recommendations. For combinatorial studies, cells were transfected with two different siRNA pools; control cells were transfected with twice as much siNT pool. 24 hours after transfection, media was changed. After 2 days, samples were collected for qPCR, and after 5 days, separate samples had cell viability determined via CellTiter-Glo (Promega, Catalog No. G7571) following manufacturer’s instructions, normalizing viability of each sample to siNT/siNT viability for each cell line.

#### Dox-inducible FRS2 and FGFR1 knockdown

shRNAs targeting FRS2 (CGCTATGGCTATGACTCGAAT) and FGFR1 (TGCCACCTGGAGCATCATAAT) were cloned into a doxycycline (dox)-inducible shRNA expression vector (Addgene, #21915). PANC0203 cells were virally transduced and selected as above. Cells were seeded into 96-well plates in the presence or absence of 1ug/mL dox. After 5 days, cell viability was determined via CellTiter-Glo (Promega, G7571) following manufacturer’s instructions.

### Immunoblotting

Cells were harvested by scraping in PBS and pelleted at 300 x g for 5 min. Pellets were lysed in high-salt RIPA buffer with protease and phosphatase inhibitor on ice for 10 min, then clarified by centrifugation at 15,000 x g for 10 min at 4 °C. Samples were separated on 4-12% NuPAGE Bis-Tris gels (Invitrogen, NP0323BOX), and transferred to PVDF membranes via overnight wet transfer. Membranes were probed with antibodies against pFGFR (1:250; CST, #3471), pAKT (1:250; CST, #13038), AKT (1:1000; CST, #2920), pERK (1:250; CST, #5726), ERK (1:1000; CST, #4695), RAS (1:1000; CST, #67648), GAPDH (1:2500; CST, #2118), MEK1 (1:1000; CST, #2352), ERK2 (1:1000; CST, #9108), or Vinculin (1:5000; Sigma-Aldrich, V9264), followed by incubation with fluorescent secondary antibodies.

### Drug response curve assays

Seeding assays were performed on each cell line to determine appropriate seeding density for confluence after 5-7 days in the control condition.

For 2D assays, cells were then seeded into 96-well plates per seeding assay results in 200μL complete growth medium the day before compound treatment. Compounds, including zoldonrasib (MedChemExpress, HY-156819), futibatinib (MedChemExpress, HY-100818), erdafitinib (MedChemExpress, HY-18708), pemigatinib (MedChemExpress, HY-109099), lirafugratinib (RLY-4008; MedChemExpress, HY-147250), TYRA-300 (SelleckChem E4727), roblitinib (FGF401; MedChemExpress, HY-101568), MRTX1133 (MedChemExpress, HY-134813), or daraxonrasib (MedChemExpress, HY-148439) were then added in serial 3-fold dilutions as monotherapy or in combination using a Tecan. Assay plates were incubated for 5-7 days and then subjected to the CTG assay following manufacturer’s instructions. Percent viability was calculated by normalizing CTG luminescence values of compound-treated wells to vehicle-treated wells.

For organoid assays, organoids were grown in organoid media as previously described (75), modified by replacing R-spondin-1- and Wnt3A-conditioned media with recombinant human R-spondin-1 (10nM; ThermoFisher, #120-38) and Wnt surrogate-Fc (0.2nM; Kactus, WNT-HM23A). At time of seeding, organoids were filtered through a 70µm filter, counted, and seeded at 1000 cells/well into 384-well ultralow attachment plates (Revvity, 6057800) into 20µL organoid media supplemented with 10% growth-factor reduced Matrigel (VWR International, #37743-722). Assay plates were incubated for 7-10 days and then subjected to the CellTiter-Glo 3D cell viability assay (Promega, G9683) following manufacturer’s instructions.

For combinatorial studies, Bliss scores were calculated via SynergyFinder 3.0 using normalized viability results (76).

### Large scale combinatorial studies

SNU410, PANC0203, or SUIT2 cells were plated into four cohorts of 2*10^7^ cells each: vehicle (DMSO), zoldonrasib (1000nM), futibatinib (300nM), or combination (zoldonrasib 1000nM + futibatinib 300nM). Cells were split and counted every 3-4 days, and all cells were replated into drug. Cohorts were maintained in this way until doubling starting cell count or until no remaining cells were detected. ZR cell lines were derived from PANC0203 and SUIT2 cells treated with zoldonrasib once these cells surpassed the starting cell count. PANC0203-ZR and SUIT2-ZR were then continuously maintained in zoldonrasib 1000nM.

### Annexin V measurement

SNU410, PANC0203, or SUIT2 cells were seeded into 96-well plates in 100μL complete growth medium at roughly 20% confluence. The next day, Incucyte Annexin V Red Dye (Sartorius, #4641) was reconstituted in PBS and diluted 1:200 into all wells. Zoldonrasib (MedChemExpress, HY-156819) and/or futibatinib (MedChemExpress, HY-100818) were then added using a Tecan. Plates were transferred to an Incucyte Live-Cell Analysis System (Sartorius), and phase contrast and red fluorescence images were acquired at 10x magnification every 2 hours. Annexin V signaling is reported as red object confluence normalized to phase object confluence within each well.

### Mouse xenograft studies

5*10^6^ SNU410 or PANC0203 cells were subcutaneously implanted in the right flank of 6-8 week-old female NU/J mice (Jackson Laboratory, strain 002019). Once tumors reached an average volume of ∼200mm^3^, mice were randomized into four groups: vehicle (5% DMSO, 40% PEG300, 5% Tween 80, 50% ddH2O), zoldonrasib (100mg/kg), futibatinib (25mg/kg), or combination (zoldonrasib 100mg/kg + futibatinib 25mg/kg), all administered via daily oral gavage. Treatment continued for 3-4 weeks, followed by holding of all further treatment. Tumor size was measured by calipers twice weekly, and tumor volume was calculated as V = 0.5 × (L × W^2^) where L = the greatest longitudinal diameter and W = the greatest transverse diameter.

### Immunohistochemistry

For hematoxylin and eosin staining, 5 µm sections were stained using a standard manual protocol. Sections were deparaffinized in xylene, rehydrated through graded alcohols to water, and stained with hematoxylin solution, Gill No. 2 (Sigma-Aldrich, GHS2128), followed by bluing reagent (Epredia, 6769001) and counterstaining with eosin Y (VWR International, 10143-132).

For pERK staining, formalin-fixed, paraffin-embedded tissue was sectioned at 5 µm onto charged slides. Staining was performed on a Leica BOND RX automated stainer (Leica Biosystems) using the Bond Polymer Refine Detection kit (DS9800) with 3,3’-diaminobenzidine (DAB) chromogen and hematoxylin counterstain. Slides were dewaxed and rehydrated on-board (Bond Dewax Solution, AR9222), and heat-induced epitope retrieval was performed for 20 min at 100 °C with Bond Epitope Retrieval Solution 2 (ER2; AR9640). pERK primary antibody (1:800; CST, #4370) was diluted in Bond Primary Antibody Diluent (AR9352) and incubated for 30 min at room temperature.

For necrosis and tumor cellularity analysis, digitized whole-slide images were analyzed in QuPath v.0.7.0. Tumor bed regions were manually annotated by a pathologist. To quantify necrosis, a random forest pixel classifier was trained on representative annotations of necrotic and viable tissue and applied to delineate necrotic areas within the tumor bed. Nuclei within the annotated tumor bed were segmented using the InstanSeg deep-learning model (QuPath InstanSeg extension v0.1.7; brightfield_nuclei-0.1.1 model) (77). A random forest object classifier was then trained on manually labeled examples to distinguish tumor from non-tumor nuclei and applied to identify tumor nuclei across the tumor bed. Necrotic area, tumor bed area, and tumor nuclei counts were exported for downstream analysis.

### Phospho-RTK analysis

PANC0203 and SNU410 cells were treated with zoldonrasib (MedChemExpress, HY-156819) for 0 or 10 days, with medium and compound refreshed every 3-4 days. At each timepoint, cells were harvested and processed per phospho-RTK array kit instructions (RCD Systems, ARY001B). Chemiluminescent signal was captured on a Bio-Rad ChemiDoc Imaging System. Spots were mapped using the provided reference spots, and spot signal intensity was quantified in Image Lab software (Bio-Rad). Phospho-RTK array spots were considered positive if having background-corrected integrated signal ≥ 10,000 units, and phospho-RTK levels are plotted for a given cell line if positive RTK signal changed by ≥ 2-fold.

### Single-cell RNA-sequencing meta-analysis

We assembled five published human PDAC droplet-based scRNA-seq cohorts (53–57). For four of these datasets (Peng, Steele, Lin, Raghavan), we used the uniformly reprocessed UMI count matrices and cell-type annotations curated by the Curated Cancer Cell Atlas (3CA) (78). For Werba (GSE205013), we obtained the raw count matrices and processed them de novo. To match 3CA annotation for the Werba dataset, we implemented a two-metric copy-number approach. Cells were filtered (≥500 genes; genes in ≥20 cells), normalized, and assigned a coarse lineage. Copy-number variation was inferred with infercnvpy on epithelial and immune/endothelial reference cells, restricted to autosomes and ordered by GENCODE v44 gene coordinates. For each sample, a tumor CNV profile was defined as the mean profile of the top 10% of epithelial cells by CNV burden. An epithelial cell was called malignant if it exceeded the 95th percentile of the reference distribution on both genome-wide CNV burden and the correlation of its CNV profile to the sample tumor profile.

We integrated the cohorts on the intersection of genes measured in all cohorts (14,939 genes). Data were normalized, reduced to 2,000 variable genes, scaled, and projected onto 50 principal components; batch effects across cohorts were corrected with Harmony (79). A neighbor graph was computed on the Harmony embedding, followed by UMAP and Leiden clustering. Gene-expression feature maps were generated by recomputing normalized expression from raw counts and mapping values onto the integrated embedding by cell barcode. For each patient and compartment, a pseudobulk profile was formed by summing UMI counts across all cells of that patient-compartment group, collapsing replicate biopsies from the same donor. Only patients with ≥10 cells in both compartments were included in statistical comparisons of CAF versus PDAC expression.

### CAF cell line derivation

Primary CAF cell lines were derived from de-identified patient tumors under an approved IRB exemption using an outgrowth method as previously described (80). Tumors were minced into pieces no larger than 1mm^3^ and cultured in Advanced DMEM/F-12 (Gibco #12634010) supplemented with 15% fetal bovine serum (FBS) (Sigma #F2442-500ML), 1x glutamine (Gibco #35050079), 1x HEPES (Corning #25-060-CI).

### CAF CM and FGF supplementation assays

CAF CM was generated by replacing the media on a confluent plate of TERT-immortalized CAFs with media without serum. After three days, CAF CM was collected and filtered through 0.45 μm filters. For CAF CM dose-response curves, cells were seeded into 96-well plates in 100μL growth medium with reduced (2%) serum. The next day, 100μL CAF CM with 2% serum or 100μL standard growth medium with 2% serum was added to cells, followed by treatment with zoldonrasib (MedChemExpress, HY-156819) and/or futibatinib (MedChemExpress, HY-100818) using a Tecan. Assay plates were incubated for 5-7 days and then subjected to the CTG assay following manufacturer’s instructions. Percent viability was calculated by normalizing CTG luminescence values of compound-treated wells to vehicle-treated wells.

For FGF supplementation dose-response curves, cells were seeded into 96-well plates in 200μL growth medium with reduced (2%) serum. The next day, heparin (5μg/mL; MedChemExpress, HY-17567A) was added to all wells, as well as DMSO, FGF1 (50ng/mL; MedChemExpress, HY-P7001), or FGF2 (50ng/mL; MedChemExpress, HY-P7004), followed by treatment with zoldonrasib (MedChemExpress, HY-156819) using a Tecan. Ligands were added at the same concentration daily for 7 days and then subjected to the CTG assay following manufacturer’s instructions. Percent viability was calculated by normalizing CTG luminescence values of compound-treated wells to vehicle-treated wells.

## ACKNOWLEDGEMENTS

Funding: This work was supported by Hirshberg Foundation grant HF-2025-118 and the Helen Hay Whitney Foundation (R.Y.E.), and by the Hale Family Center for Pancreatic Cancer Research, the Kerr Family Fund for Translational Research, and the WHH Foundation (J.M.C.).

## Author contributions

R.Y.E. and W.R.S. conceived the project, wrote the manuscript, and provided project leadership. R.Y.E., Y.H., J.J., D.T.F., and E.E. built the digenic screening library. R.Y.E., A.Z., T.T., and A.A. performed screens. Y.H., G.L., and A.D., developed screen analysis methodology and provided bioinformatics support. R.Y.E., A.Z., E.K., and V.Y. conducted validation experiments. I.C.M., C.U.J, N.N.K.V., S.A.Z., C.L., and J.J.Y. developed CAF models, analyzed CAF samples, and provided CAF experimental support. K.E.D., N.Y., and Y.Y.T. provided organoid experimental support. A.J.A. and J.M.C. provided clinical and translational guidance.

## Conflict of Interest

R.Y.E. has received consulting fees from Third Rock Ventures and Luma Group, which are not related to this work. J.M.C. receives research funding to his institution from Amgen, Artios, Partners Therapeutics, Roche, Servier, Synnovation, and Bristol-Myers Squibb. He receives research support from Bayer, Caris, GSK, Arcus Biosciences, and Pyxis; he has also received honoraria from Partners Therapeutics, BeOne, Prelude, Abdera, Gilead, and Elevar Therapeutics, AstraZeneca, Bayer, and Akesobio. W.R.S. is or was a board or scientific advisory board member and/or holds equity in Atavistik Bio, CJ Bioscience, Delphia Therapeutics, Ideaya, Pierre Fabre, RedRidge Bio, Epidarex Capital, 2Seventy Bio, and Scorpion Therapeutics; has consulted for Array, Astex, Ipsen, Merck, Sanofi, and Servier; and receives or received research funding from Bayer Pharmaceutical, Bristol-Myers Squibb, Boehringer-Ingelheim, Ideaya, Calico, Novartis, Servier, Pfizer, Merck, Ridgeline Discovery, and BridgeBio.

## Data and materials availability

CRISPR screen LFC and Combinatory score values generated in this study are provided as Supplementary Data. All custom analysis code will be made publicly available on GitHub.

