## Supplementary Figures, and Supplementary Table legends for "Context-specific genetic interaction mapping reveals combinatorial KRAS/FGFR dependence in pancreatic cancer"

### Supplementary Figure 1

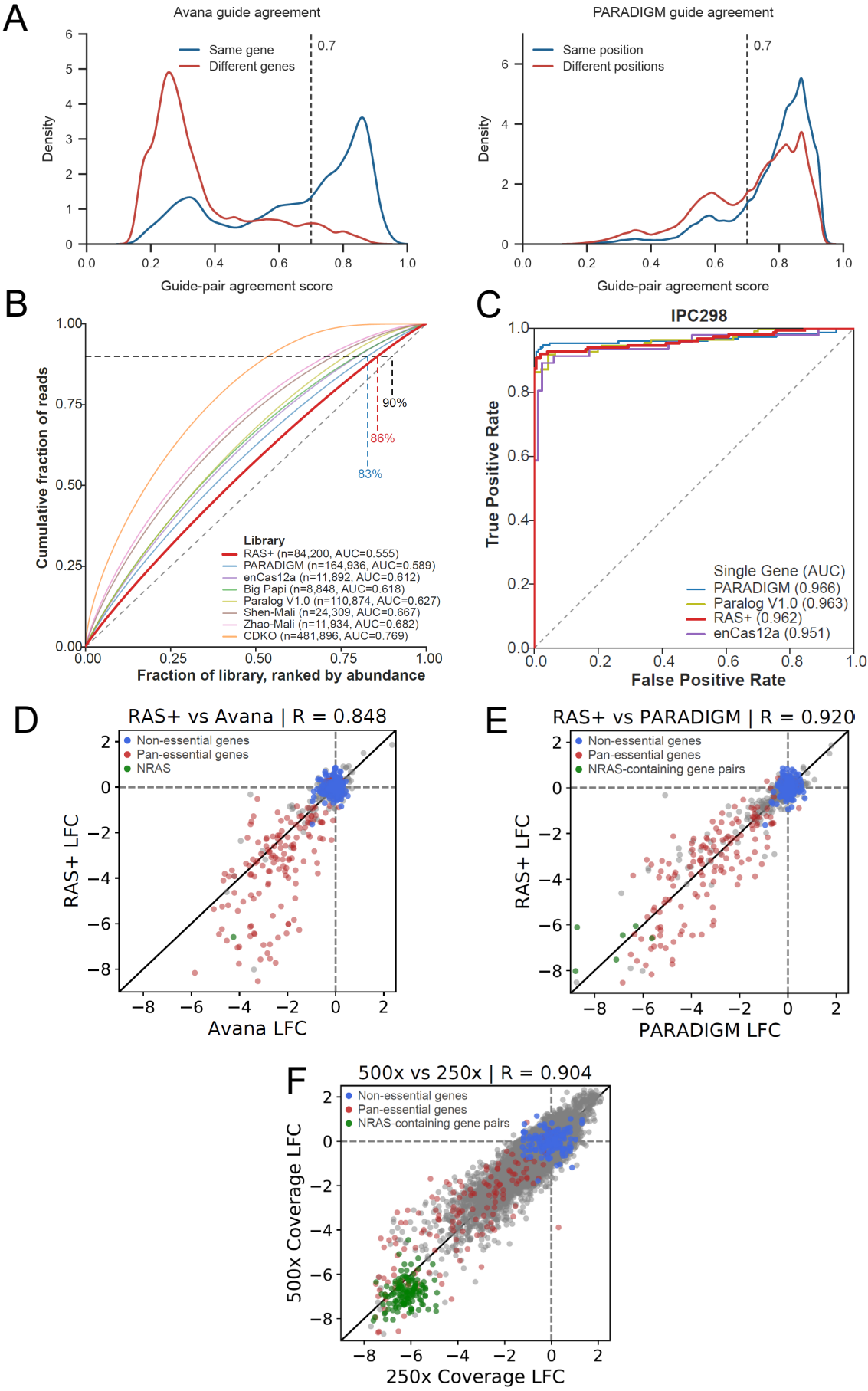

##### **Supplementary Figure 1. RAS+ library metrics.**

(A) Distributions of guide pair agreement scores generated by a random forest model trained on Avana screening data. Left, scores for guide pairs targeting the same gene or different genes in the Avana training set. Right, scores for guide pairs targeting the same gene in PARADIGM screens, with the gene-targeting sgRNAs occupying the same or different positions in the digenic vector. Dashed lines indicate the threshold used to prioritize guide pairs (agreement score > 0.7). (B) Area-under-the-curve (AUC) analysis for library distribution from various digenic CRISPR screening libraries. (C) Single-gene receiver operating characteristic (ROC)-AUC curve derived from pan-essential and non-essential genes in IPC298. (D) LFC of genes shared between RAS+ and Avana screening in IPC298. (E) LFC of gene pairs shared between RAS+ and PARADIGM screening in IPC298. (F) LFC of gene pairs after RAS+ screening at 250x or 500x coverage. In D-F, non-essential (blue) and pan-essential (red) individual genes are highlighted, as well as gene pairs containing NRAS (green).

Supplementary Figure 2

A

| Cell line | Lineage | RAS+ mutation(s) |
| --- | --- | --- |
| BXPC3 | PDAC | BRAF V487_P492delinsA |
| COLO205 | CRC | BRAF V600E |
| COLO678 | CRC | KRAS G12D |
| HCT15 | CRC | KRAS G13D; PIK3CA E545K, D549N; ERBB3 N126K |
| HPAFII | PDAC | KRAS G12D |
| IPC298 | Melanoma | NRAS Q61L |
| KP363T | CRC | BRAF V600E; PIK3CA E545K |
| PACADD165 | PDAC | No RAS+ mutations |
| PANC0203 | PDAC | KRAS G12D |
| PANC1 | PDAC | KRAS G12D |
| SNU410 | PDAC | KRAS G12D |
| SNU478 | Ampullary | No RAS+ mutations |

B

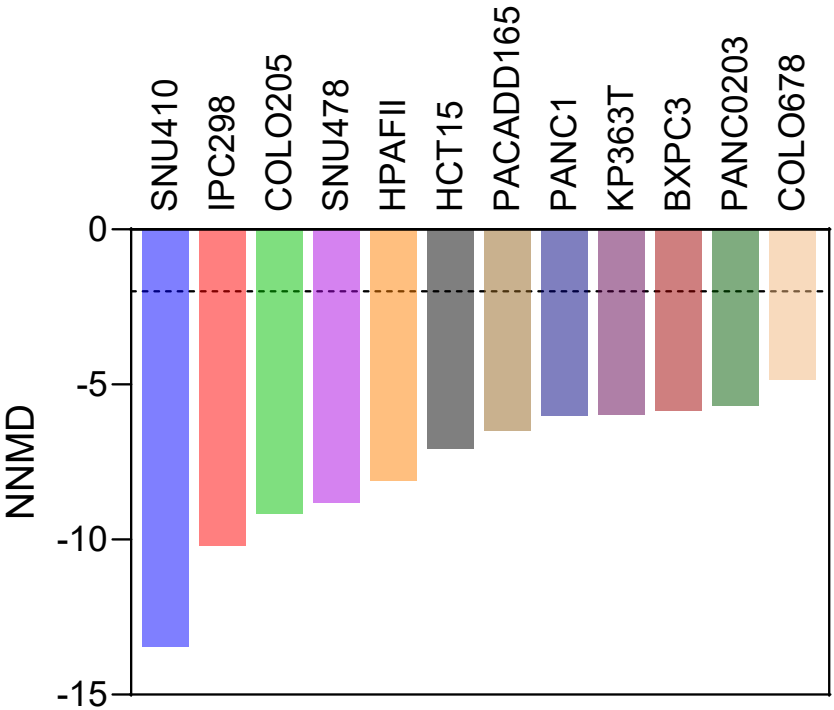

**Supplementary Figure 2. 12 RAS+ screen metrics.**

(A) Table of cell lines screened with the RAS+ library, along with lineage and driver mutations in RAS+ genes. (B) Null-normalized mean difference (NNMD) scores for lines screened with the RAS+ library. Screens with NNMD values $\leq -2$ are considered to pass QC evaluation.

### Supplementary Figure 3

**BXPC3**
BRAT V487\_P492delinsA
PDAC

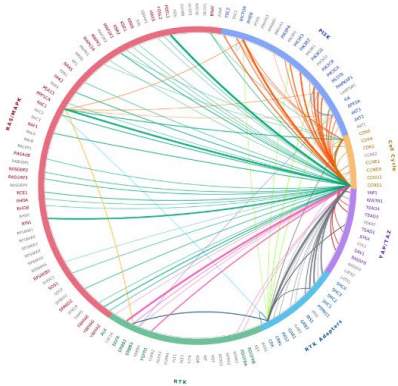

**COLO205**
BRAF V600E
CRC

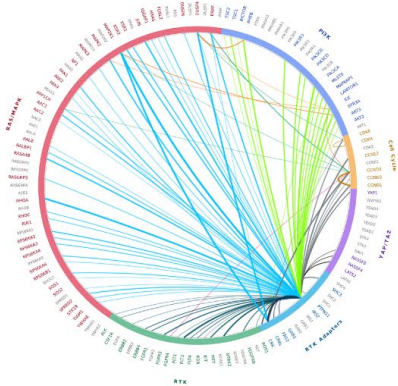

**COLO678**
KRAS G12D
CRC

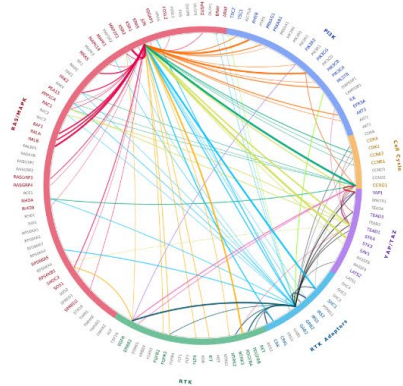

**HCT15**
KRAS G13D; PIK3CA E545K, D549N; ERBB3 N126K
CRC

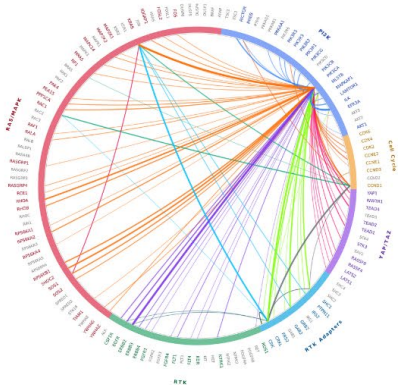

**HPAFII**
KRAS G12D
PDAC

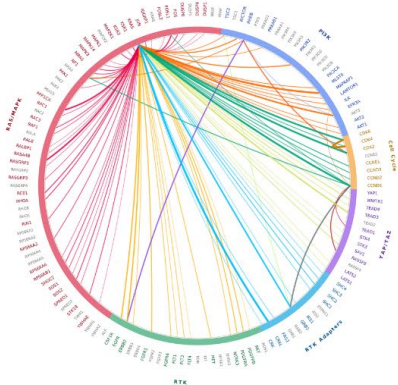

**IPC298**
NRAS Q61L
Melanoma

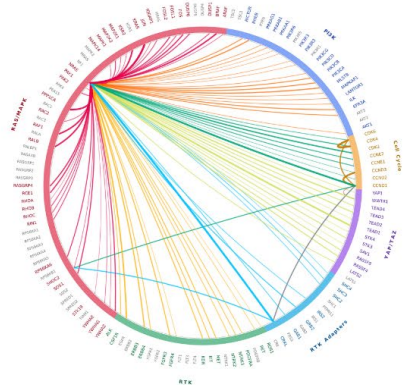

**KP363T**
BRAF V600E; PIK3CA E545K
CRC

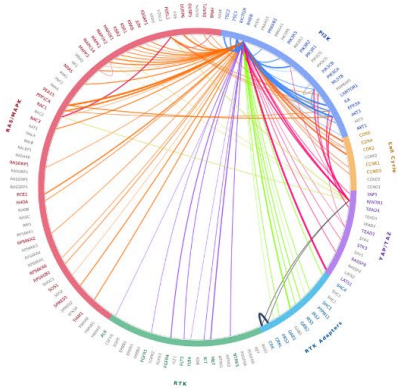

**PACADD165**
no RAS+ mutations
PDAC

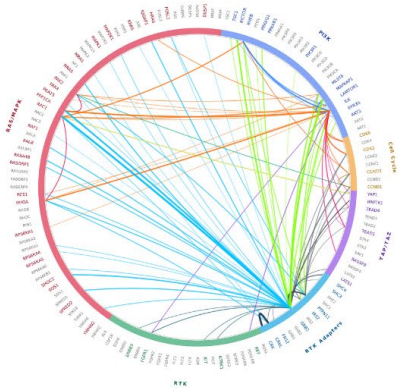

**PANC1**
KRAS G12D
PDAC

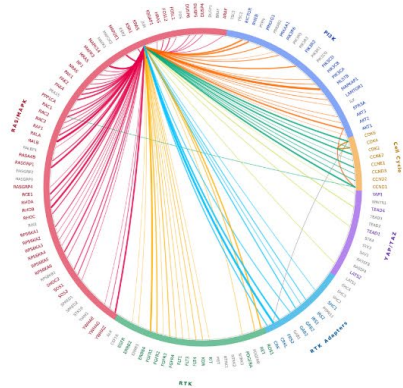

**PANC0203**
KRAS G12D
PDAC

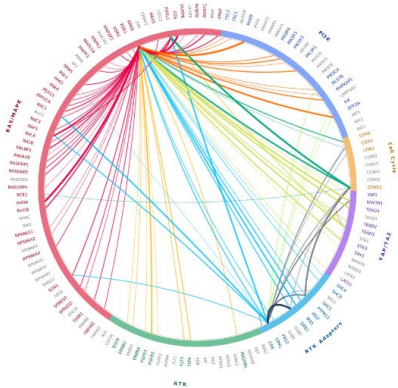

**SNU410**
KRAS G12D
PDAC

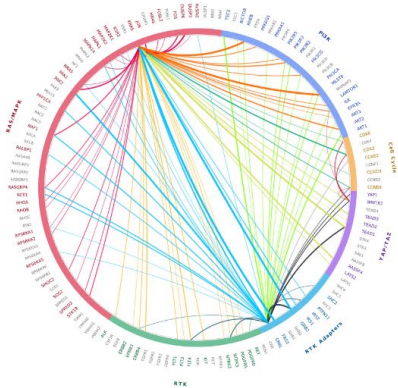

**SNU478**
no RAS+ mutations
BTC

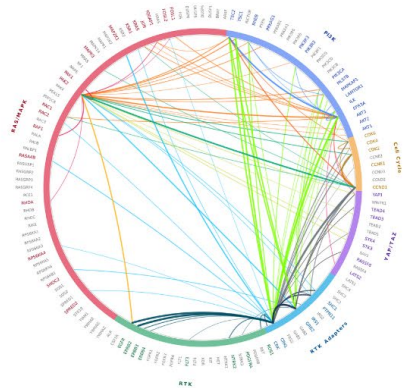

**Supplementary Figure 3. 12 RAS+ screen lethality maps.**

Lethality maps showing the 100 most lethal gene pairs in each cell line screened with the RAS+ library. Chords are colored by which gene categories are connected by each chord, and greater width indicates greater lethality.

### Supplementary Figure 4

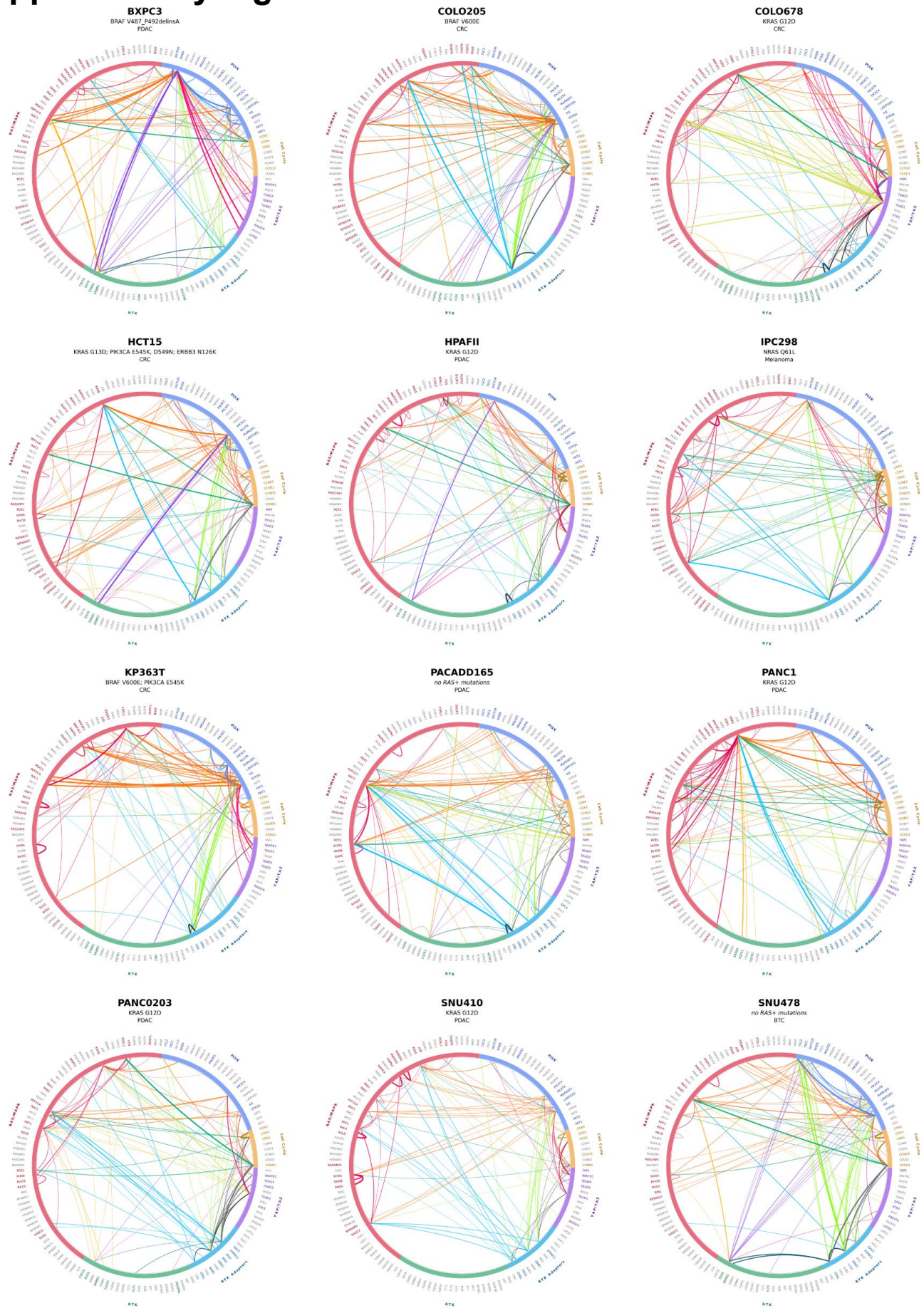

**Supplementary Figure 4. 12 RAS+ screen synergy maps.**

Synergy maps showing the 100 most synergistic gene pairs in each cell line screened with the RAS+ library, as defined by lowest Combinatory scores. Chords are colored by which gene categories are connected by each chord, and greater width indicates greater synergy.

Supplementary Figure 5

A

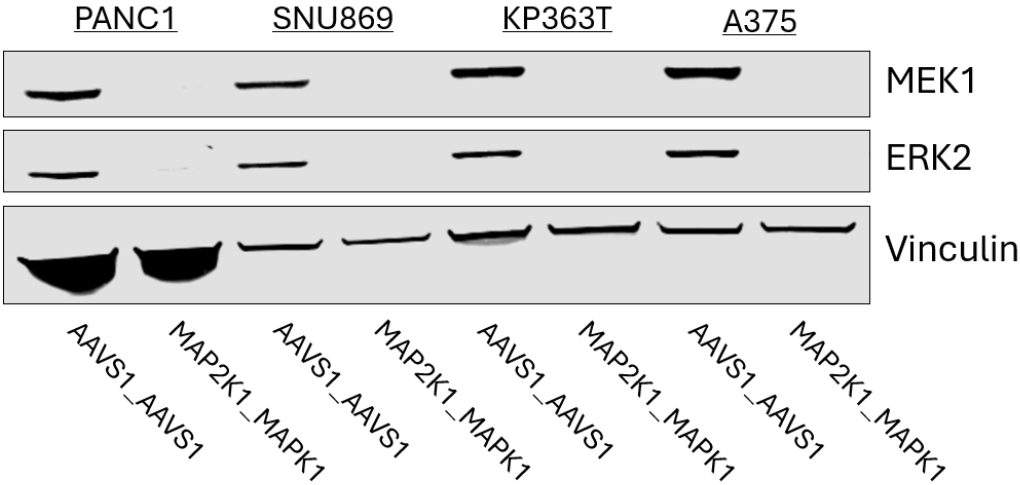

B

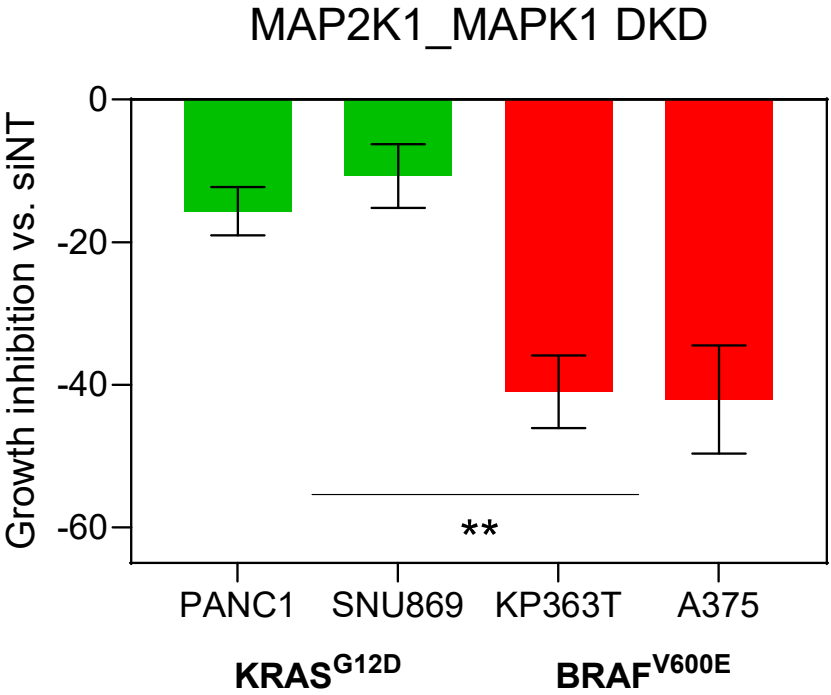

**Supplementary Figure 5. MEK1/ERK2 is a digenic dependency in BRAF<sup>V600E</sup> but not KRAS<sup>mut</sup> lines.**

(A) Immunoblots for MEK1 and ERK2 of PANC1, SNU869, KP363T, and A375 cells with dual expression of sgRNAs against MAP2K1 and MAPK1 (MAP2K1\_MAPK1) or control (AAVS1\_AAVS1). (B) siRNA dual knockdown of MAP2K1 and MAPK1 leads to significantly reduced growth in BRAF<sup>V600E</sup> vs. KRAS<sup>mut</sup> lines. Viability measured by CellTiter-Glo and compared to viability of cells exposed to siNT. Data represent mean ± SD. *P* value calculated by two-tailed unpaired Student's *t* test. \*\*: *P* < 0.01.

Supplementary Figure 6

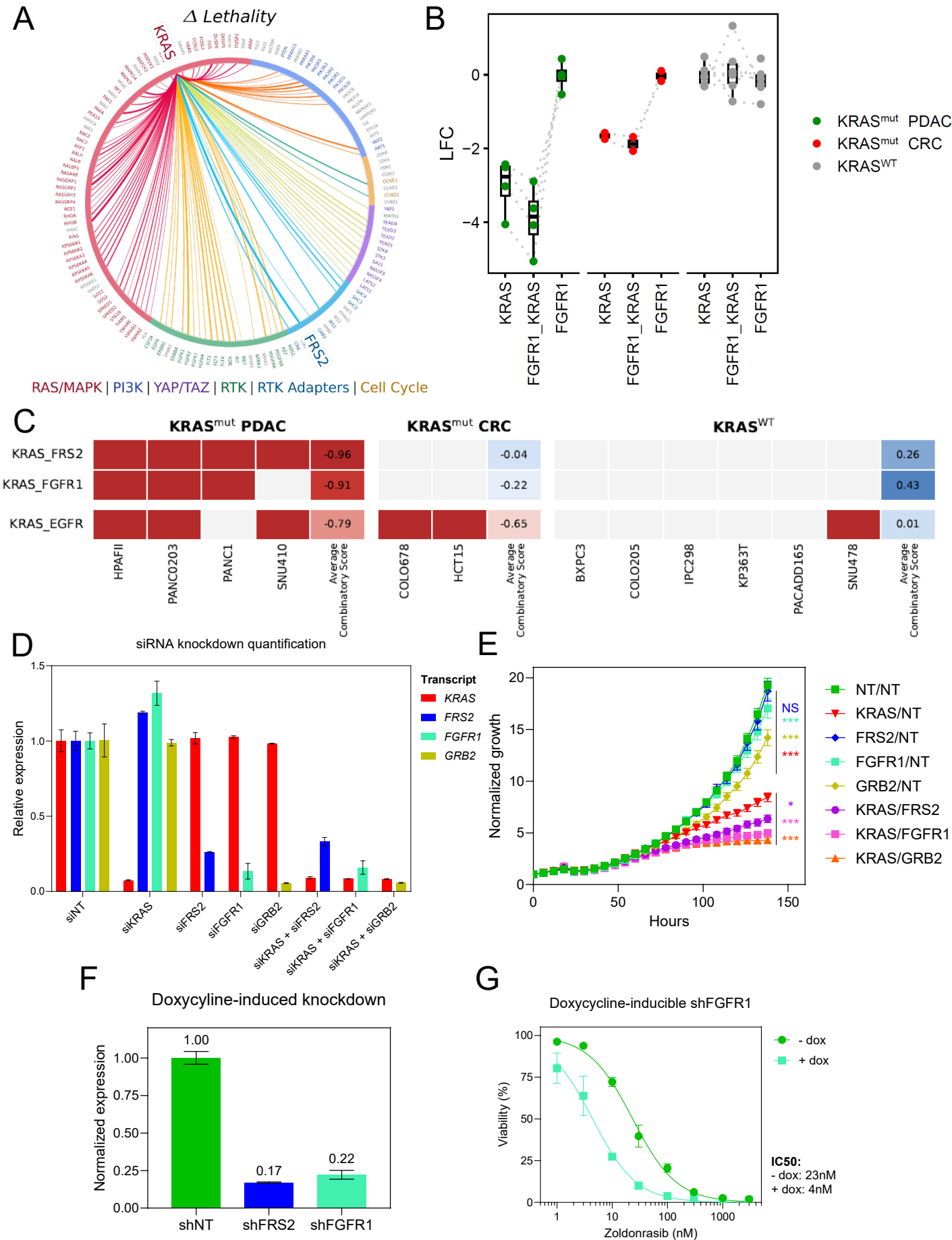

**Supplementary Figure 6. KRAS/FRS2 is the strongest differential dependency in KRAS<sup>mut</sup> PDAC.**

(A) Lethality map showing the 100 most differentially lethal gene pairs in KRAS<sup>mut</sup> PDAC lines versus all others. Chords are colored by which gene categories are connected by each chord, and greater width indicates greater lethality. KRAS\_FRS2 is the most differentially lethal gene pair. (B) Boxplot displaying LFC scores for KRAS\_FGFR1 and corresponding single genes, separated into KRAS<sup>mut</sup> PDAC, KRAS<sup>mut</sup> CRC, and KRAS<sup>WT</sup> lines. (C) Combinatory scores for KRAS\_FRS2, KRAS\_FGFR1, and KRAS\_EGFR across all lines screened with the RAS+ library. Combinatory scores $\leq -0.50$ indicated in red. (D) qPCR quantification of siRNA single or dual knockdown in SNU410 cells, corresponding to Fig. 3F. Data are normalized to mean expression in cells with siNT. Data represent mean ± SD. (E) siRNA dual knockdown of KRAS and FGFR-pathway genes leads to significantly reduced growth versus single knockdown in PANC0203 cells. Cell growth was assayed using Incucyte and normalized to timepoint 0. Data represent mean ± SD. *P* values calculated by extra sum-of-squares *F* test comparing growth rate constants of exponential growth curve fits. Single gene knockdowns were compared versus siNT/siNT, and dual gene knockdowns were compared versus siKRAS/siNT. \*\*\*: *P* < 0.001, \*: *P* < 0.05, NS: *P* > 0.05. (F) qPCR quantification of shRNA knockdown after doxycycline-induction in PANC0203 cells, corresponding to Figs. 3G and S6G. Data are normalized to mean expression in cells with siNT. Data represent mean ± SD. (G) Zoldonrasib dose-response curve of PANC0203 cells with or without doxycycline induction of an shRNA targeting FGFR1. Viability measured by CellTiter-Glo. Data represent mean ± SD.

### Supplementary Figure 7

A

Zoldonrasib/FGFRi

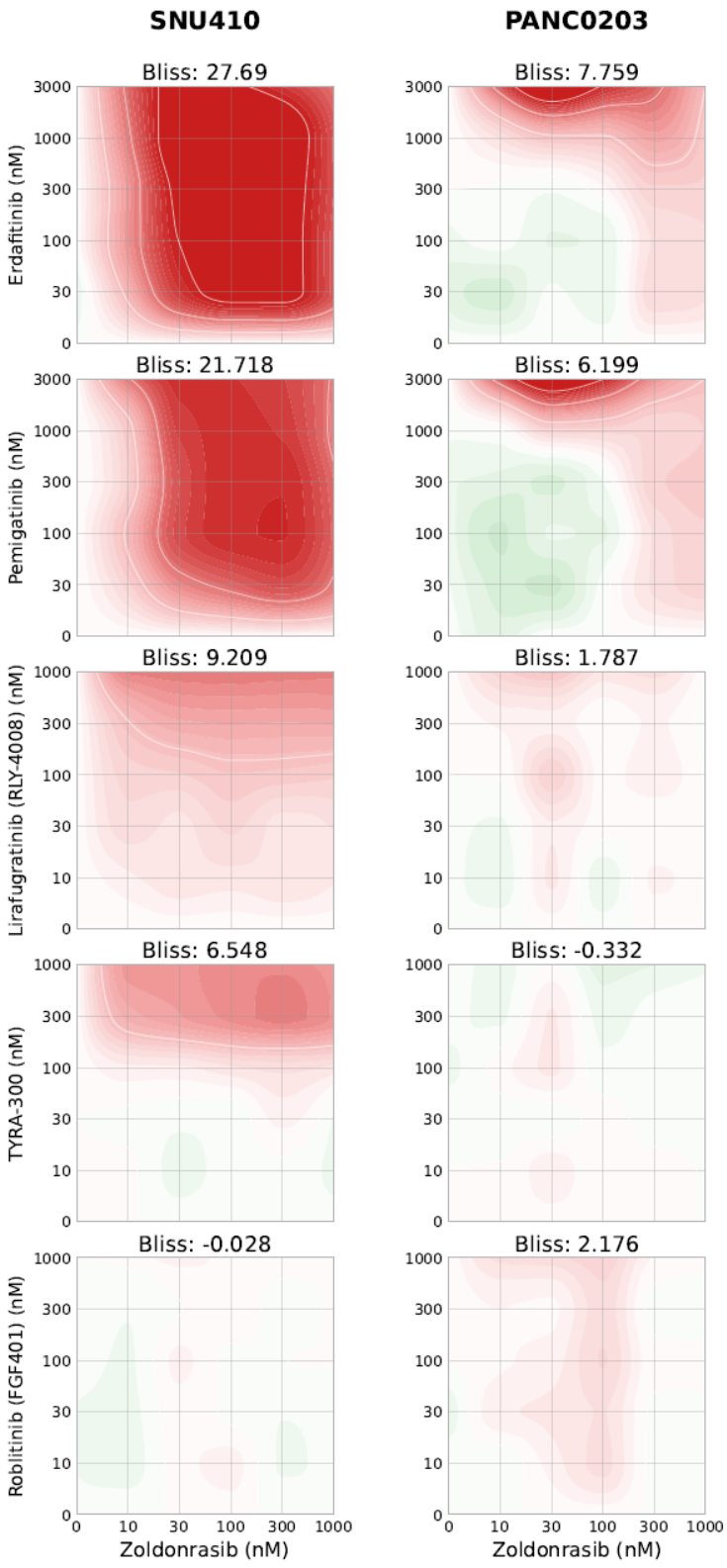

B

RASi/Futibatinib

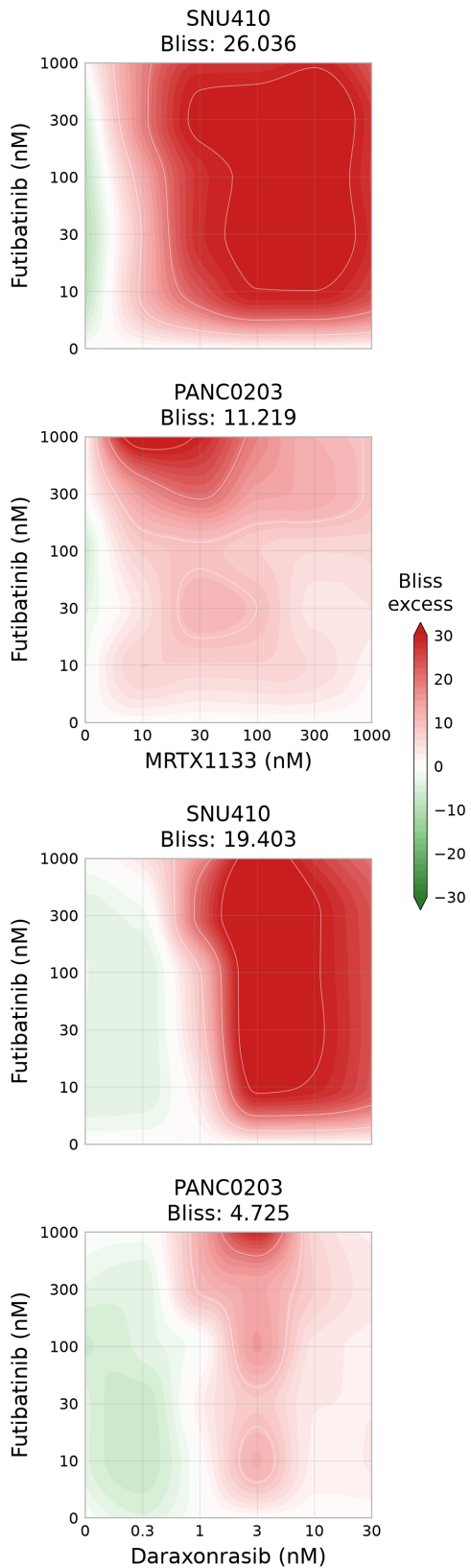

**Supplementary Figure 7. KRAS and FGFR inhibitors demonstrate synergy in KRAS<sup>mut</sup> PDAC.**

(A) Bliss synergy matrices for SNU410 and PANC0203, testing the combination of zoldonrasib with various multi-FGFR and paralog-selective FGFR inhibitors. (B) Bliss synergy matrices for SNU410 and PANC0203, testing the combination of futibatinib with the KRAS<sup>G12D</sup> inhibitor MRTX1133 or the pan-RAS inhibitor daraxonrasib. For all plots, viability was measured by CellTiter-Glo, and Bliss synergy scores were calculated across the full dose-response matrix.

Supplementary Figure 8

A

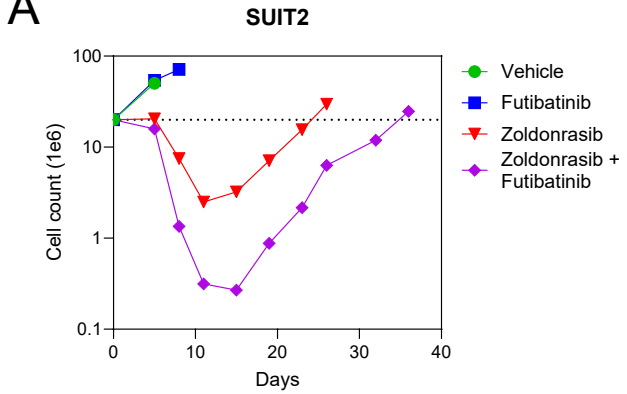

B

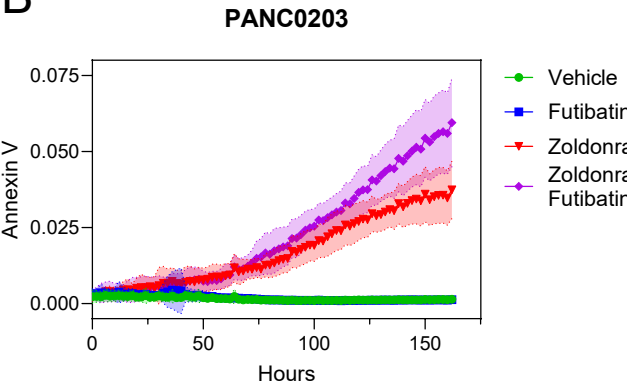

C

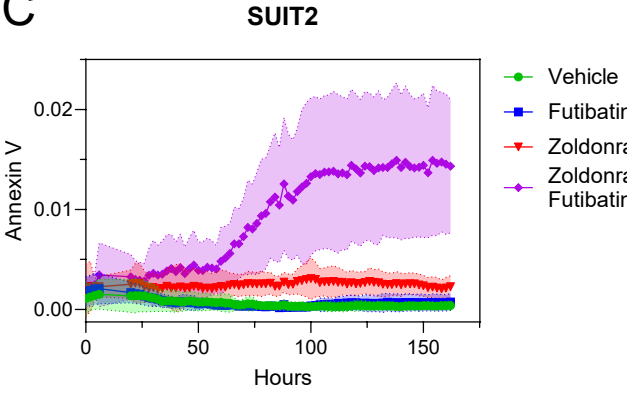

D

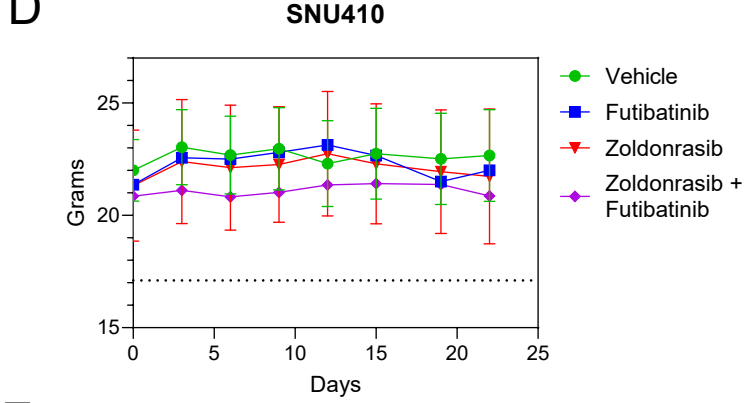

E

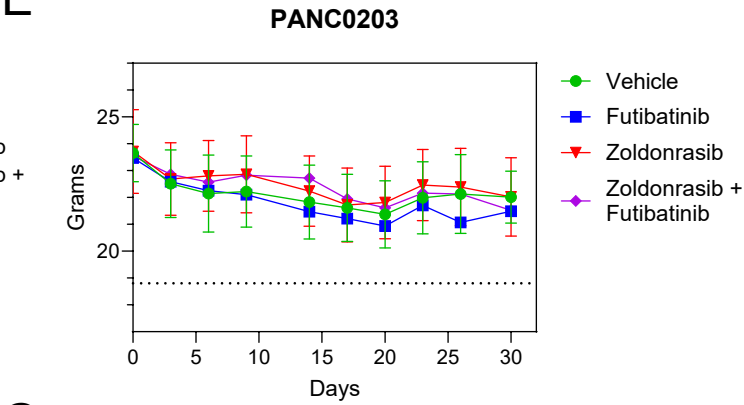

G

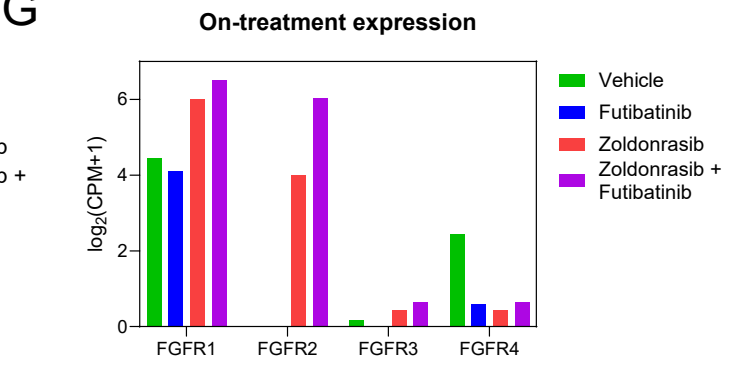

F

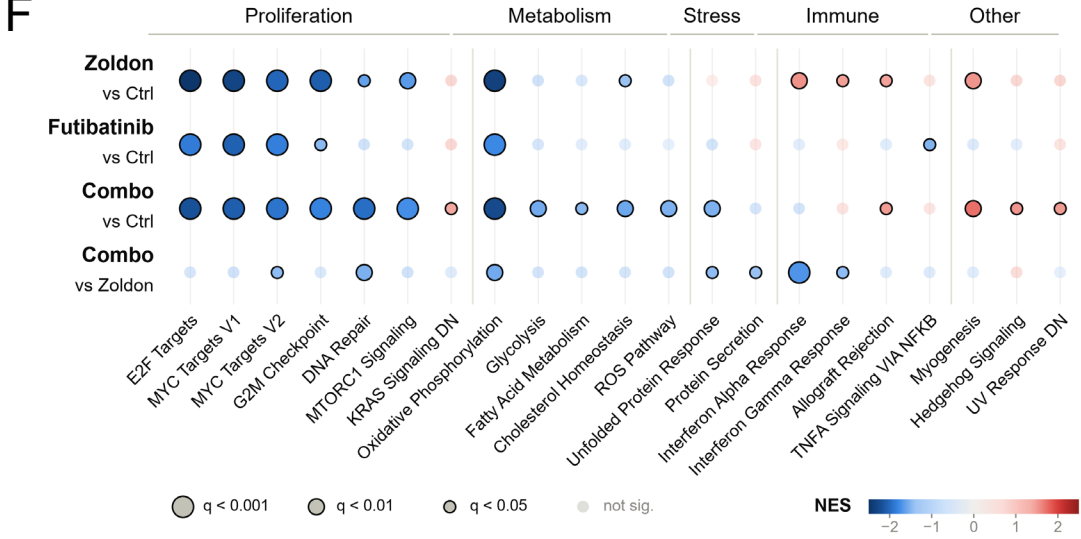

**Supplementary Figure 8. Zoldonrasib/futibatinib is highly cytotoxic against KRAS<sup>mut</sup> PDAC.**

(A) Large scale, longitudinal drug treatment experiments in SUIT2, treated with zoldonrasib 1000nM, futibatinib 300nM, combination, or vehicle. Cell counts are plotted over time on a log<sub>10</sub>-scaled y-axis. The dotted line indicates the starting cell number of $20 \times 10^6$ cells. (B) Annexin V staining of PANC0203 cells treated with zoldonrasib 100nM, futibatinib 1000nM, combination, or vehicle. Annexin V staining was assayed using Incucyte and normalized to confluence. (C) Annexin V staining of SUIT2 cells treated with zoldonrasib 300nM, futibatinib 1000nM, combination, or vehicle. Annexin V staining was assayed using Incucyte and normalized to confluence. (D-E) Weight of mice with SNU410 (D) or PANC0203 (E) tumors receiving daily zoldonrasib 100mg/kg, futibatinib 25mg/kg, combination, or vehicle, corresponding to Fig. 4E. (F) GSEA for Hallmarks of Cancer gene set (81) enrichment in comparisons of zoldonrasib versus vehicle, futibatinib versus vehicle, combination versus vehicle, and combination versus zoldonrasib-treated PANC0203 tumors. All gene sets with $q < 0.05$ in any comparison are shown. NES: normalized enrichment score. (G) Bar plot showing mean $\log_2(\text{CPM}+1)$ for FGFR1-4 in PANC0203 tumors receiving zoldonrasib, futibatinib, combination, or vehicle.

### Supplementary Figure 9

A

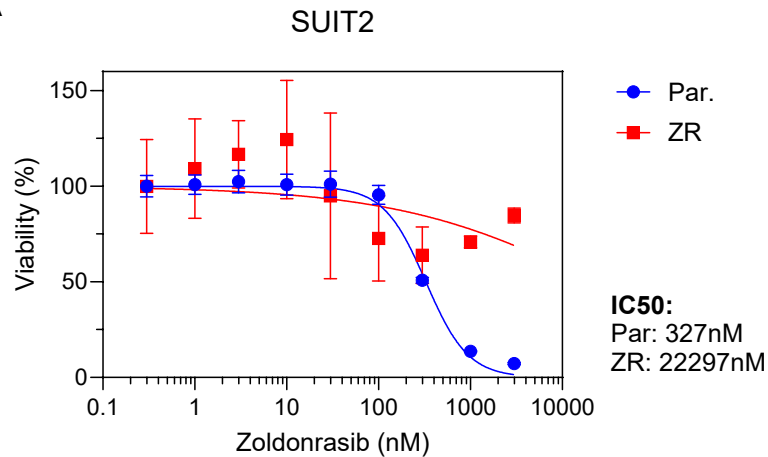

B

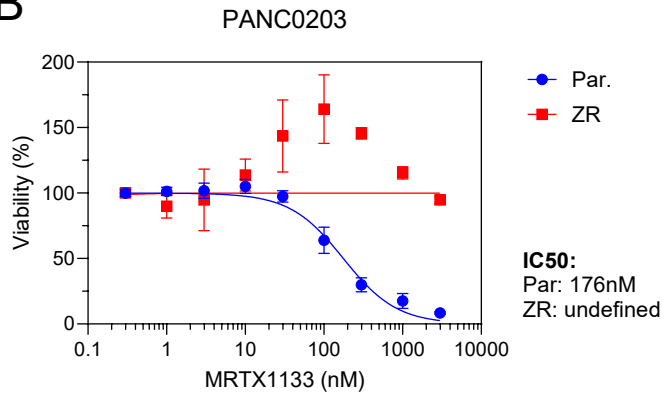

C

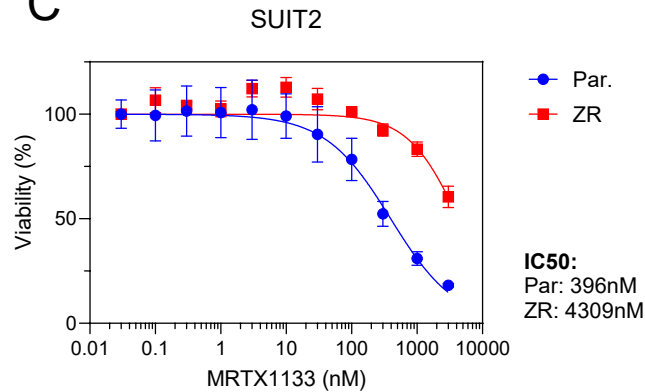

D

E

F

**Supplementary Figure 9. Prolonged KRAS inhibition leads to adaptive FGFR signaling.**

(A) Zoldonrasib dose-response curve of SUIT2 parental versus zoldonrasib-resistant (ZR) cells. (B-C) MRTX1133 dose-response curves of PANC0203 (B) or SUIT2 (C) parental versus ZR cells. (D-E) Daraxonrasib dose-response curves of PANC0203 (D) or SUIT2 (E) parental versus ZR cells. Viability for all dose-response curves measured by CellTiter-Glo. Data represent mean ± SD. (F) Bar plot showing mean voom-normalized log2 CPM for FGFR1-4 in SUIT2 parental and ZR cells. Data represent mean expression across biological replicates. *P* values calculated using empirical Bayes moderated *t* test. \*\*\*: *P* < 0.001.

Supplementary Figure 10

**Supplementary Figure 10. An integrated PDAC patient scRNA-seq atlas identifies CAFs as the primary source of FGF ligands in patient PDAC tumors.**

(A-B) UMAP of 364,082 cells from 97 patients across five PDAC scRNA-seq cohorts (53–57), colored study of origin (A) or by patient of origin (B). (C) Dot plot of FGF ligands and FGFRs in the PDAC or CAF compartment, pooled across all five scRNA-seq cohorts and split by primary and metastatic disease. Dot size denotes the percentage of cells in the compartment expressing the gene; dot color denotes average compartment expression. (D) Analysis of individual FGFR paralog detection at the patient level among the 97 patients in the meta-analysis. Receptors were scored as detected if expressed in at least 1%, 5%, or 10% of a patient's malignant cell compartment.

### Supplementary Figure 11

## A

## B

## C

**Supplementary Figure 11. CAFs are the primary source of FGF ligands from in vitro models.**

(A) Expression of all 18 secreted FGF ligands in 17 CAF versus 13 PDAC organoid models. *Q* values calculated via Welch's two-sample *t*-test followed by Benjamini-Hochberg correction. *Q* listed as ND for ligands with median CPM <1 in both compartments. (B) Expression of ligands reaching significance in (A), restricted to patients for whom both a CAF and organoid line were derived. *Q* values calculated via paired *t*-test followed by Benjamini-Hochberg correction. Lines connect models derived from the same patient. (C) Expression of FGFR1-4. *Q* values calculated via Welch's two-sample *t*-test followed by Benjamini-Hochberg correction. For all plots, dotted line represents CPM = 1.

Supplementary Figure 12

A

B

C

**Supplementary Figure 12. PDAC CAFs promote KRAS inhibitor resistance via paracrine activation of FGFR signaling.**

(A) Zoldonrasib dose-response curves of PANC0203 (left) and SUIT2 (right) supplemented with or without CAF CM. Viability measured by CellTiter-Glo. Data represent mean ± SD. (B) Immunoblots for pFGFR, pERK, and pAKT of SNU410 (left), PANC0203 (middle), and SUI2 (right) with or without 4 hours of CAF CM exposure. (C) Growth inhibition of zoldonrasib 100nM, futibatinib 1000nM, combination, or vehicle in PANC0203 (left) and SUIT2 (right) supplemented with or without CAF CM. Viability measured by CellTiter-Glo. Data represent mean ± SD.

### Supplementary Figure 13

**Supplementary Figure 13. Individual FGF ligands promote KRAS inhibitor resistance.**

(A) Zoldonrasib dose-response curves of SUIT2 with supplementation of FGF1, FGF2, or vehicle. Viability measured by CellTiter-Glo. Data represent mean ± SD. (B-C) Zoldonrasib dose-response curve of SNU410 (B) or SUIT2 (C) with lentiviral overexpression of GFP, FGF1, or FGF2. Viability measured by CellTiter-Glo. Data represent mean ± SD. (D) Zoldonrasib x futibatinib Bliss synergy matrices for SNU410 with lentiviral overexpression of FGF2. Viability was measured by CellTiter-Glo, and Bliss synergy score was calculated across the full dose-response matrix.

#### **TABLE LEGENDS**

##### **Supplementary Table 1. RAS+ library genes.**

List of the 144 genes assessed in all pairwise combinations in the RAS+ library and associated gene categorization.

##### **Supplementary Table 2. RAS+ library constructs.**

List of the 84,200 digenic constructs in the RAS+ library, with associated sgRNA sequence, sgRNA source, and construct type.

##### **Supplementary Table 3. 12 screen LFC table.**

Gene pair LFC for 12 RAS+ screens.

##### **Supplementary Table 4. 12 screen Combinatory score table.**

Gene pair Combinatory scores for 12 RAS+ screens.

##### **Supplementary Table 5. In vivo GSEA.**

GSEA for Hallmarks of Cancer gene set enrichment in comparisons of zoldonrasib-, futibatinib-, combination-, or vehicle-treated PANC0203 tumors.

##### **Supplementary Table 6. Phospho-RTK arrays after KRAS inhibition.**

Background-corrected integrated pixel intensity for phospho-RTK array spots for PANC0203 and SNU410 after 0 or 10 days of zoldonrasib 300nM.

##### **Supplementary Table 7. FGF/FGFR expression in CAFs and organoids.**

CPM of all secreted FGFs, FGFR1-4, and FRS2 in a panel of 17 PDAC CAFs and 13 PDAC organoids.
